# Predicting biological activity from biosynthetic gene clusters using neural networks

**DOI:** 10.1101/2024.06.20.599829

**Authors:** Hemant Goyat, Dalwinder Singh, Sunaina Paliyal, Shrikant Mantri

**Author notes:** **Corresponding authors:** Dalwinder Singh, Shrikant Mantri. These authors contributed equally (Hemant Goyat, Dalwinder Singh).

## Abstract

Microorganisms like bacteria and fungi have been used for natural products that translate to drugs. However, assessing the bioactivity of extract from culture to identify novel natural molecules remains a strenuous process due to the cumbersome order of production, purification, and assaying. Thus, extensive genome mining of microbiomes is underway to identify biosynthetic gene clusters or BGCs that can be profiled as particular natural products, and computational methods have been developed to address this problem using machine learning. However, existing tools are ineffective due to a small training dataset, dependence on old genome mining tools, lack of relevant genomic descriptors, and prevalent class imbalance. This work presents a new tool, NPBdetect, that can detect multiple bioactivities and has been designed through rigorous experiments. Firstly, we composed a larger training set using MIBiG database and a test set through literature mining to build and assess the model respectively. Secondly, the latest antiSMASH genome mining tool was used to obtain BGC and introduced new sequence-based descriptors. Thirdly, neural networks are used to build the model by dealing with class imbalance issues through the class weighting technique. Finally, we compared the NPBdetect tool with an existing tool to show its efficacy and real-world utility in detecting several bioactivities with high confidence.

## 1 Introduction

A biosynthetic gene cluster (BGC) is a group of two or more genes within a genome that are physically close to each other. These genes collaboratively encode the biosynthetic pathway responsible for producing a specialized metabolite and its chemical variants [1]. These specialized metabolites, also known as secondary metabolites or natural products, are not only directly involved in growth and reproduction but also provide the producing organism with selective advantages like helping in metal transport, symbiotic association, and acting as competitive agents against other microbes by displaying bioactivities such as antibiotic, antifungal, siderophore and many more [2]. Thus, these natural products have substantial applications in agriculture, clinical, and pharmaceutical industries as well as therapeutics [3].

Natural products have gained the attention of the scientific community due to their diverse use. However, finding molecules with useful functions is like finding a needle in a haystack. From the natural source of bacteria, thousands of strains can be accumulated during microbial isolation. To get the desired function or bioactivity from the extract of a bacterial culture, pure bacterial isolates are obtained from the primary ones, and previously probed microorganisms are excluded from screening [4]. Afterwards, assaying, production, and purification processes are used to identify compounds which are eventually screened for their bioactivity. These experimental bottlenecks not only make the identification of novel natural products difficult owing to cumbersome processes but are also associated with time, cost, and bioactivity hit rates [4]. To address the ongoing antibiotic resistance crisis, these approaches including efficient ones like mass spectrometry [5], NMR [5] and MicroED [6, 7] are not enough [8].

One promising approach to address this issue is the prediction of the bioactivity of molecules using their BGCs [9, 10]. Various genome mining tools like antiSMASH [11], Deep-BGC [12], and PRISM [13] have been developed to identify BGCs from the genome. Also, some tools were developed to detect classes of natural products. It includes PKminer [14] for discovering Polyketide biosynthetic class, RiPPMiner [15], RODEO [16], NeuRiPP [17], and DeepRipp [18] for mining ribosomally synthesized and post-translationally modified peptides (RiPPs) and BAGEL [19] for identifying putative bacteriocin ORFs (antimicrobial peptides). Consequently, these tools mined vast amounts of genomic data and found millions of BGC sequences. For instance, BiGFAM database contains over 1.2 million BGCs [8]. To predict bioactivity based on BGC, a handful of tools are available. DeepBGC is one such tool that utilizes a machine learning (ML) approach to predict antibacterial, antifungal, cytotoxic, and inhibitor bioactivities. A small training set of 370 instances was used, which makes it less reliable for new predictions. Another BGC-based tool is the Natural Product Function (NPF) [20] which also employed an ML approach and extracted features from 14 BGC descriptors to detect six bioactivities. This tool used a relatively large training set for BGC detection and feature extraction. Three classifiers, extra-randomized trees (ERTree), support vector machine (SVM), and regularized linear models with stochastic gradient descent learning (SGD) classifier, were used to build models by optimizing their parameters using 10-fold cross-validation (CV) and balanced accuracy. The ML models were developed independently for each bioactivity, resulting in 18 models that increase the risk of under and overfitting. Recently, NPF tool was extended to the detection of fungal secondary metabolites. This NPFF tool [21] is trained on NPF dataset along with 314 new fungal BGCs.

Even though outstanding tools have been developed for bioactivity detection, we have identified several shortcomings in addressing this problem more effectively. It includes overfitted models due to fine-tuning classifiers to an individual classification problem while relying on a small dataset, lack of assessment for BGC descriptors, reliance on older versions of antiSMASH to obtain feature information, and utilizing few bioactivities. This work complements the ongoing efforts to address the issue of bioactivity prediction with artificial intelligence (AI). We proposed a new tool, NPBdetect, to detect multiple bioactivities. It is built through rigorous experiments using a larger training set. Firstly, this work improves data standardization by composing two datasets, one training and one test set which is inspired by contemporary datasets in AI. To facilitate consistent and systematic deposition and retrieval of data on BGC, Minimum Information about a Biosynthetic Gene Cluster (MIBiG) database [1] is utilized. The training set comprises the MIBiG BGCs where bioactivity labels were available in either MIBiG database or NPF dataset [20]. The test set, on the other hand, is independent and composed through an extensive literature review to uncover the bioactivities of compounds synthesized by BGCs. Secondly, we have proposed new sequence-based descriptors that have been proven to perform better than other descriptors when combined with PFAM domains. Lastly, eight bioactivities have been selected to build the model where class-imbalance is prevalent. The proposed model not only outperformed the existing approach but also detects siderophore bioactivity with high accuracy, thereby validating its efficacy for practical use.

## 2 Materials and Methods

### 2.1 Data description

AI approaches in the natural products domain are still in their infancy and suffer from a lack of high-quality datasets [9]. Although data preparation is labor-consuming, high-quality data is vital for model generalization [22]. To develop the NPBdetect tool, new datasets have been compiled and the multi-label classification problem has been extended by incorporating five more bioactivities, namely, siderophore, antiprotozoal, inhibitor, antiviral, and surfactant, besides the previously attempted antibacterial, antifungal and cytotoxic/antitumor bioactivities.

Figure 1 presents the data curation process for NPB training and test datasets. We proposed one training and one test dataset to evaluate the ML models, as shown in Figure 1A. The new training set consists of 1295 BGCs which are selected by reviewing NPF training set and adding new entries from MIBiG database. While developing the NPF tool, literature was curated to find the bioactivity of BGCs listed in the MIBiG 1.4 database. Then, 1152 BGCs from MIBiG were selected, and 1003 BGCs were identified with antibacterial, antifungal, antitumor or cytotoxic, antifungal or cytotoxic/antitumor, anti-gram positive, and anti-gram negative bioactivities. However, NPF tool used all BGCs for feature extraction even though label information is available for 1003 instances only. So, we named it NPF-filtered dataset which is also used to build, evaluate, and compare models. To further refine the NPF dataset with up-to-date information, we validate the remaining instances in MIBiG 3.1 database. We found 56 BGCs where label information is no longer available and removed them to obtain NPF-latest dataset. This dataset is used to select the initial set of BGCs in NPB training set. Additionally, the recent MIBiG database has 42 more bioactivities. Thus, we have also included those activities where labels for more than 40 BGCs are available. Finally, eight classes from 1295 high-confidence BGCs are selected to build the model. On the other hand, test set is composed through literature mining only and named NPB-LM. The details about this dataset and its analysis are provided further (subsection **NPB-LM test dataset**).

**Figure 1:**
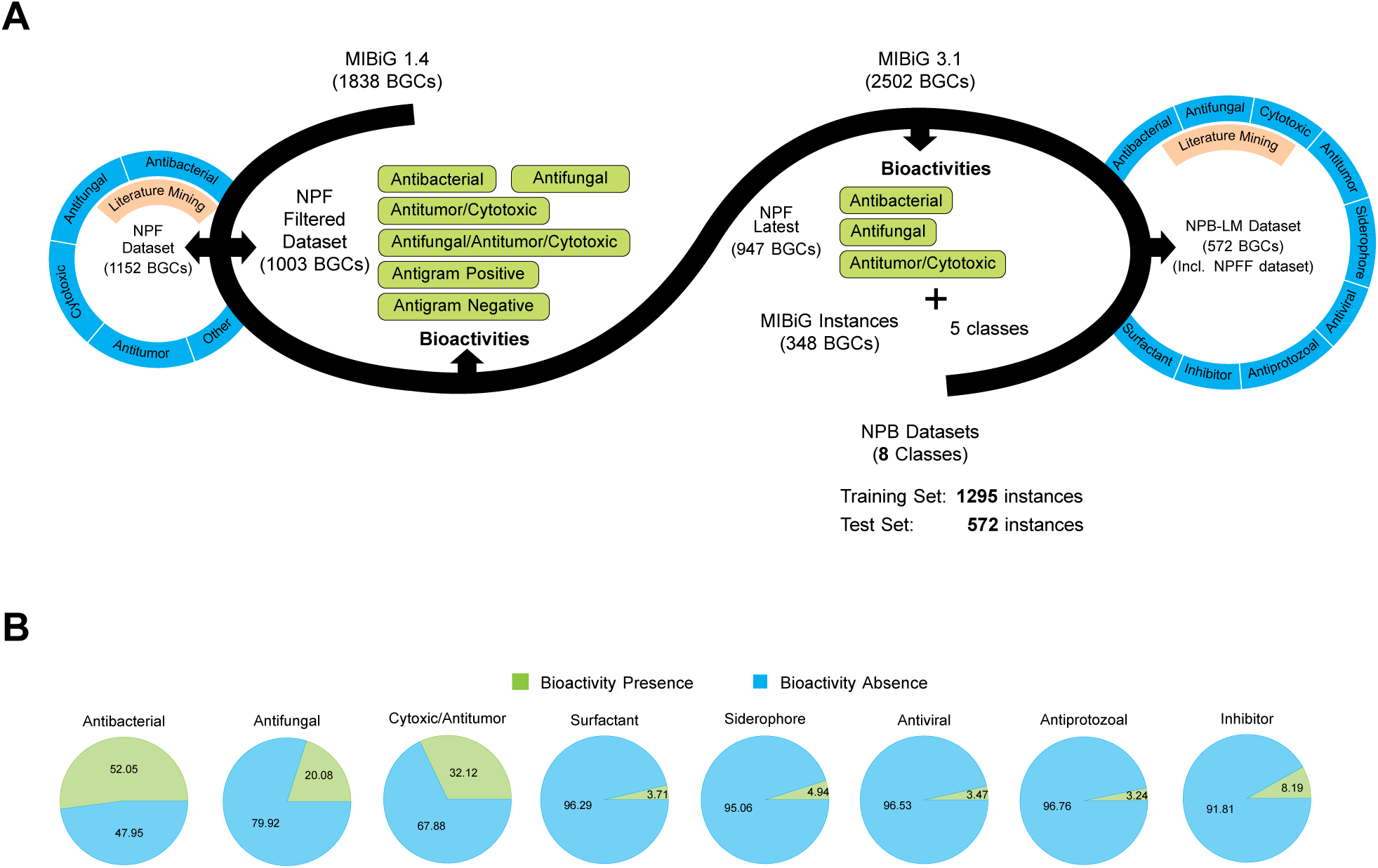
(A) Curation of NPB dataset using MIBiG, NPF, and NPFF datasets, and through literature review. (B) Class imbalance of bioactivities in NPB training set.

Figure 1B presents the class imbalance of bioactivities in the training set. The class imbalance is prevalent in this classification problem because of the limited information available for most bioactivities. Only antibacterial class is balanced, whereas antifungal and cytotoxic/antitumor classes have a reasonable imbalance. The remaining siderophore, antiprotozoal, inhibitor, antiviral, and surfactant classes are highly imbalanced which makes this classification problem challenging.

### 2.2 NPB-LM test dataset

Even though the latest MIBiG database is expanding with large-scale validation and re-annotation of existing entries, bioactivity information is available for less than half of the BGCs. So, plenty of BGCs have undocumented bioactivity, and through a comparative analysis of the MIBiG database and NPF dataset, we identified 1207 such BGCs. Out of these BGCs, 517 BGCs with the desired biological activities are found. For the remaining BGCs without any documented bioactivity, overlap with NPFF dataset containing 314 fungal BGCs is performed. We found 55 fungal BGCs with the desired biological activities from this dataset. In total, 572 BGCs have bioactivites, and their distribution is shown in Figure 2A. The most prominent group of dataset encodes antibacterial (36.5%), followed by cytotoxic/antitumor (17.8%) and antifungal (16.0%) compounds. For the newer bioactivities, we found fewer validations (*<* 10%) in the literature. Figure 2B presents the relative frequency between NPB-LM and MIBiG database BGCs. The bioactivity information was found for many MIBiG entries, particularly for anticancer/antitumor and siderophore categories. The full details of literature mining in terms of BGC bioactivity and corresponding publication(s) are provided in Supplementary Table S1.

**Figure 2:**
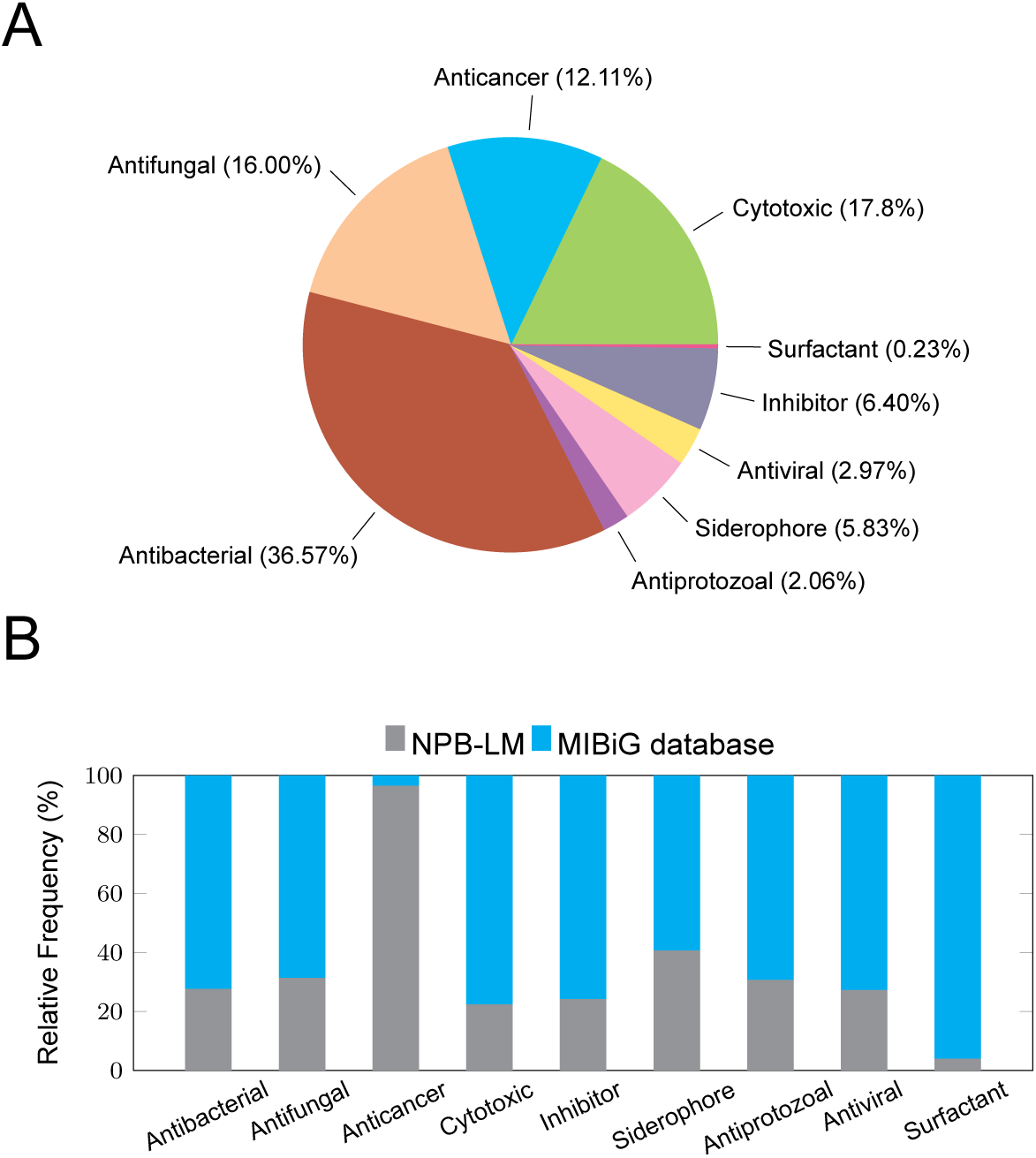
(A) Percentage of bioactivity distribution for NPB-LM BGCs. (B) Illustration of relative frequency between NPB-LM and MIBiG BGCs for various bioactivities

We have categorized BGCs into distinct biosynthetic classes as shown in Table 1. We also compared the MIBiG database (1123 BGCs) previously annotated with bioactivity in MIBiG 3.1 and the newly annotated NPB-LM dataset. Remarkably, we observed a consistent pattern in both sets, with the highest number of BGCs falling into the Polyketide biosynthetic class, followed by Non-Ribosomal Peptide (NRP) BGCs. Also, bioactivity distribution within these classes exhibited a similar trend, with antibacterial BGCs being the most prevalent, followed by cytotoxic BGCs.

**Table 1:** distribution of biosynthetic classes in NPB-LM dataset.

| BGC Class | NPB-LM | Bioactivities |  |  |  |  |  |  |  |
| --- | --- | --- | --- | --- | --- | --- | --- | --- | --- |
|  |  | Antibacterial | Antifungal | Cytotoxic | Antiprotozoal | Antiviral | Siderophore | Surfactant | Inhibitor |
| Polyketide | 155 | 86 | 54 | 89 | 6 | 7 | 0 | 1 | 18 |
| NRP | 128 | 67 | 26 | 39 | 4 | 6 | 25 | 0 | 11 |
| NRP, Polyketide | 79 | 30 | 18 | 39 | 3 | 4 | 12 | 0 | 9 |
| RiPP | 51 | 44 | 0 | 7 | 1 | 1 | 0 | 0 | 4 |
| Terpene | 44 | 21 | 13 | 23 | 1 | 0 | 0 | 0 | 5 |
| Other | 69 | 40 | 20 | 28 | 0 | 7 | 14 | 1 | 4 |
| Alkaloid | 21 | 12 | 5 | 12 | 1 | 0 | 0 | 0 | 4 |
| Saccharide | 5 | 5 | 0 | 0 | 0 | 0 | 0 | 0 | 0 |
| Terpene, Polyketide | 4 | 2 | 0 | 4 | 1 | 0 | 0 | 0 | 1 |
| Saccharide, Other | 2 | 2 | 0 | 0 | 0 | 0 | 0 | 0 | 0 |
| NRP, Saccharide | 2 | 2 | 0 | 2 | 0 | 0 | 0 | 0 | 0 |
| Polyketide, Saccharide | 2 | 2 | 0 | 2 | 0 | 0 | 0 | 0 | 0 |
| Polyketide, NRP, Saccharide | 2 | 2 | 2 | 0 | 0 | 1 | 0 | 0 | 0 |
| Alkaloid, Polyketide | 3 | 2 | 1 | 3 | 0 | 0 | 0 | 0 | 0 |
| NRP, Other | 2 | 1 | 0 | 2 | 1 | 0 | 0 | 0 | 0 |
| Other, Polyketide | 1 | 0 | 0 | 1 | 0 | 0 | 0 | 0 | 0 |
| Alkaloid, Terpene, Saccharide | 1 | 1 | 1 | 1 | 0 | 0 | 0 | 0 | 0 |
| Polyketide, NRP, Other | 1 | 1 | 0 | 1 | 0 | 0 | 0 | 0 | 0 |
| **Total** | **572** | **320** | **140** | **253** | **18** | **26** | **51** | **2** | **56** |

We also conducted taxonomic comparisons at the genus and phylum levels for species possessing bioactive BGCs in MIBiG database and NPB-LM dataset, as shown in Figure 3. In the MIBiG database, *Actinobacteria* is most prominent at the phylum level, accounting for 58% of bioactive entries, followed by *Proteobacteria* at 19%, and *Firmicutes* at 9% as shown in Figure 3A. However, in the NPB-LM dataset, we observed a shift in this trend. *Ascomycota* is prevalent at the phylum level, representing 37.9% of entries followed by *Ascomycota*, *Actinobacteria* at 26% and *Proteobacteria* at 15%. Notably, there were some previously unreported phylum entries, including *Streptophyta*, *Basidiomycota*, *Planctomycetes*, *Rhodophyta*, and *Bacillariophyta* as shown in Figure 3B. We compared the top 10 genera at the genus level where *Streptomyces* has the highest bioactive entries in both sets. In the MIBiG database, *Pseudomonas* and *Bacillus* followed as the next prominent genera as shown in Figure 3C. In contrast, *Aspergillus* took the second spot, followed by *Fusarium* in the NPB-LM dataset as shown in Figure 3D.

**Figure 3:**
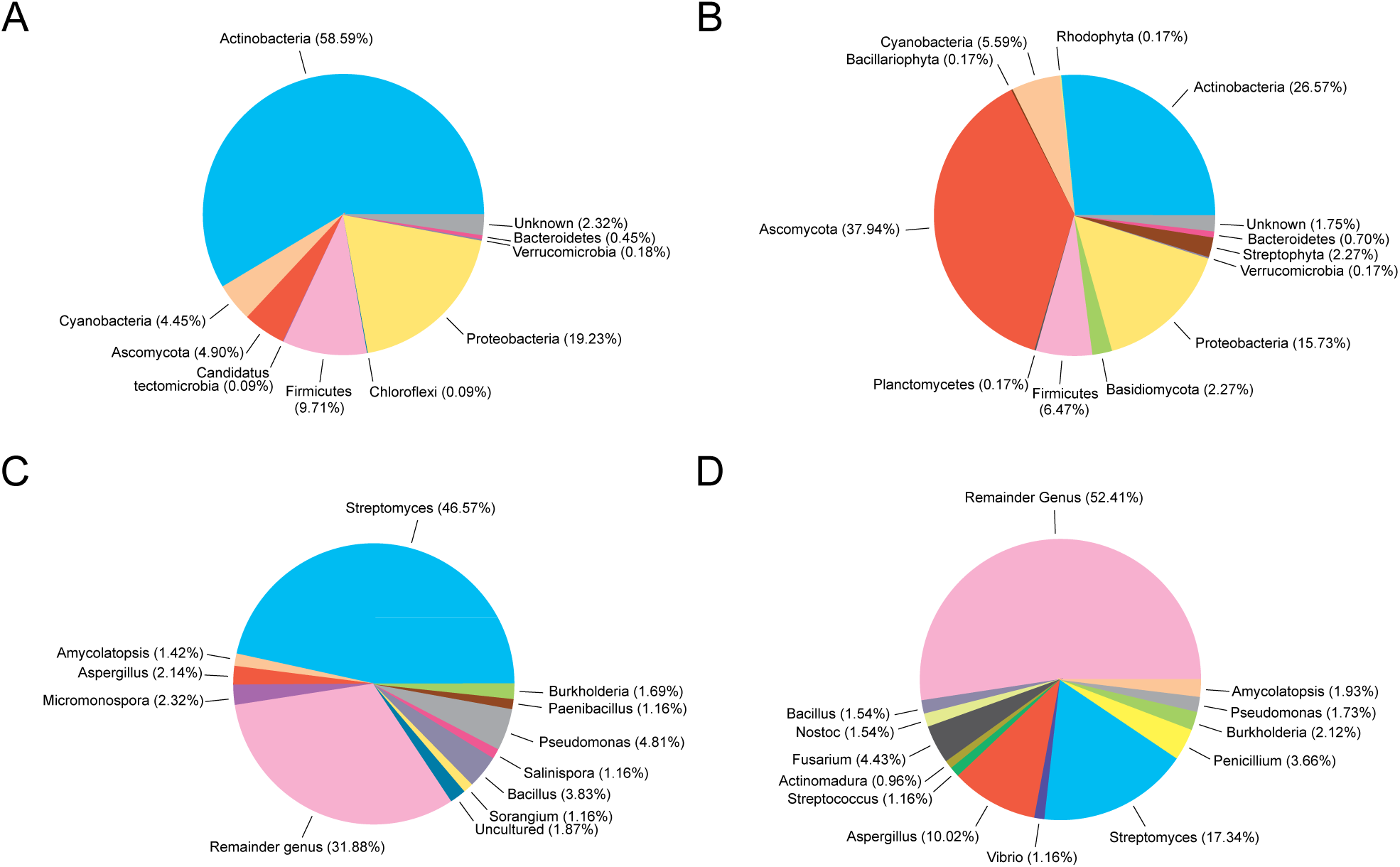
Taxonomic proportions of phylum distribution of (A) MIBiG database (1123) and (B) NPB-LM dataset (572). Genus distribution of (C) MIBiG database and (D) NPB-LM dataset

### 2.3 Methods

#### 2.3.1 Literature Mining

Various databases of the natural product domain encompass information related to natural products including gene annotations, structure, tailoring enzymes, and bioactivity. However, a few databases have comprehensive information about natural products compiled on one platform. MIBiG [23] is one such database that provides most of the information related to a natural product and is widely regarded as the benchmark for BGCs. It has been continuously updated and managed through community-driven initiatives. MIBiG 3.1 database had recorded data on the bioactivity of 1123 out of 2502 entries. Additionally, previous literature mining efforts [20] yielded bioactivity data for 1152 BGC sequences. To systematically identify unique entries for NPB training set, we eliminated the overlapping entries and obtained 1207 BGC-SM pairs that lacked recorded bioactivity information.

For the NPB test set, we have employed a literature-mining approach to gather information on the documented biological activities of each secondary metabolite. The relevant publications were sourced from PubMed and Google Scholar. For a particular BGC ID, we retrieved the compound information from MIBiG 3.1 database and used the compound name as bait to look for its biological activities. If a compound has one or multiple biological activities, then we have considered it bioactive, and its corresponding BGC IDs have been included in the dataset. When a BGC was reported to generate multiple products, we deemed it active if at least one of the products exhibited activity. If a compound has no documented biological activity and no proof of experimental validation for its inactivity, we concluded that it had not been tested for its biological activity and, therefore, was not included in our dataset.

#### 2.3.2 Feature extraction

To develop the NPBdetect tool, we relied on antiSMASH annotations, unlike different types of descriptors used in NPF tool. Specifically, PFAM domains and sequence composition of BGCs in terms of nucleotide and protein information are used for feature extraction from the annotations. The details of the feature extraction are given as follows:

##### antiSMASH annotations

As mentioned previously, antiSMASH annotations played an essential role in developing the classification models for bioactivity prediction. NPF tool used the antiSMASH 4.1 annotations for feature extraction from PFAM, secondary metabolism Clusters of Orthologous Groups (smCOG), coding sequence (CDS) motifs, and predictions of monomers for polyketides and non-ribosomal peptides descriptors. Further, features were extracted by sequence similarity networks (SSNs) of PFAM domains and Resistance Gene Identifier (RGI). From the 14 descriptors and 1152 instances, 1809 features were computed to build the models.

We have also used these annotations for feature extraction. In particular, we have used antiSMASH 7 [24] annotations for feature extraction to leverage the continuous refinement of rules to detect BGCs. We selected PFAM through experiments with descriptors from antiSMASH annotations, SSNs, and RGI. The relevancy of descriptors is measured independently and in combinations where PFAM emerged as the most relevant. So, similar to NPF tool, PFAM entries in annotation files were extracted and used as features. The features were computed at a range of bit-score and frequency thresholds to optimize the descriptor parameters.

##### Sequence Composition

The sequence composition of the genome is also helpful in determining the bioactivities because of the dependence between adjacent amino acids in proteins. Thus, the composition of the genome and proteins detected with antiSMASH are used for feature extraction. The sequence composition, particularly nucleotide content, has been widely used to classify protein-coding and non-coding RNAs [25, 26, 27]. It captures the biases in nucleotide and peptide frequencies and can be computed using several ways, such as standard and log-likelihood ratios, to measure differential usage between the presence and absence of bioactivity. We used both ratios in this work, depending on the sequence size. The features of large nucleotide sequences of BGC are computed using log-likelihood ratio whereas features of small protein sequences are calculated using standard ratio.

To capture the sequence composition, multiple *k-*mer frequencies between the presence and absence of bioactivity are computed initially from the training set. These frequencies are computed for each bioactivity independently. For a given BGC DNA sequence, the sequence probability under the presence and absence of bioactivity is calculated. Then, a score is computed using the log-likelihood ratio to measure differential usage between the occurrence of bioactivity. Specifically, a probability profile is built by computing the *k*-mer frequencies from the training datasets for presence 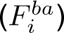 and absence 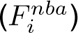 of bioactivity where *i* = 0, 1*, …,* 4*^k^*. Then, frequencies are normalized by the total counts to obtain probabilities. A score is computed for DNA sequence as follows:

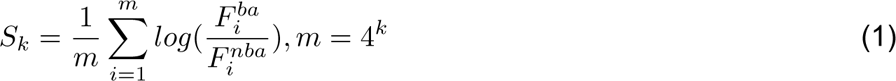

The score determines the relative degree of *k*-mer usage bias in a particular sequence where positive values indicate the presence of bioactivity. In contrast, negative values indicate an absence of bioactivity. We have experimented with 4, 6, and 8-mer profiles in this work.

Similarly, we have used protein sequence composition to design a new descriptor. BGC is composed of proteins ranging from tens to hundreds. Thus, amino acids in these proteins may also play an essential role in identifying the bioactivities. Unlike nucleotide sequences which generate one feature per sequence, the sequence composition of protein is captured independently.

Owing to the small size of proteins, the ratio between the presence and absence of bioactivity is computed. Firstly, the probability profile is built by computing the *k*-mer frequencies of all proteins from the training dataset for presence 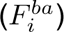 and absence 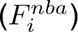 of bioactivity where *i* = 0, 1*, …,* 22*^k^* that denotes the twenty standard amino acids including two ambiguous B and Z amino acids. Afterward, frequencies are normalized by the total counts to obtain probabilities. Here, the score for each protein is computed as follows:

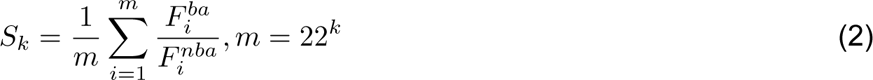

The process is repeated for maximum *N* proteins, and the obtained scores are concatenated to form a feature vector for each BGC. We have used a maximum of 50 proteins to compute the scores for each bioactivity. Thus, overall 400 features are generated for each BGC. Due to the large combinatorial space of amino acids, we have computed profiles using *k* = 1 and 2 only.

#### 2.3.3 Neural Networks

The recent advances in deep learning have drastically improved the accuracy of neural networks (NN). Over the last decade, researchers have proposed several effective techniques to make NN a state-of-the-art learning mechanism to solve complex problems and identify patterns from data [28]. It includes dropout, normalization of batches and layers, non-linear activation functions, optimizers, and loss functions [28, 29, 30]. Besides these advances, NN also supports the multi-label classification natively. Thus, NPBdetect tool is developed using the intrinsic capabilities of dense neural networks for bioactivity prediction. Figure 4 presents the multilayer neural model using PFAM and sequence-content descriptors to identify bioactivities. This feedforward neural network consists of an input layer that takes 1,311 features as input, two hidden layers to learn non-linear relationships between the input and output data, and an output layer with eight nodes to predict bioactivities. The number of neurons at the hidden layers is 100 and 50. The activation function for hidden layers is the Gaussian Error Linear Unit (GELU) [31]. The sigmoid function is used at the output layer. Additionally, layer normalization was used after each hidden layer to improve the model generalization. The model is trained using a backpropagation algorithm to optimize the network parameters while minimizing the binary cross-entropy loss function. The loss function for multi-label classification problem **x** = [*x*_1_*, x*_2_*, …, x_C_*] is given as follows:

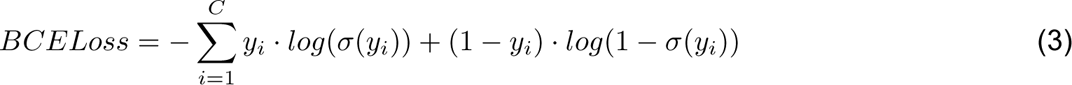

where *σ*(*·*) is the sigmoid function, *y_i_*denotes the actual label of bioactivity (Presence:1, Absence: 0), *σ*(*y_i_*) indicates the prediction probability that activity is present. *C* denotes the total classes which are 8 for our model.

**Figure 4:**
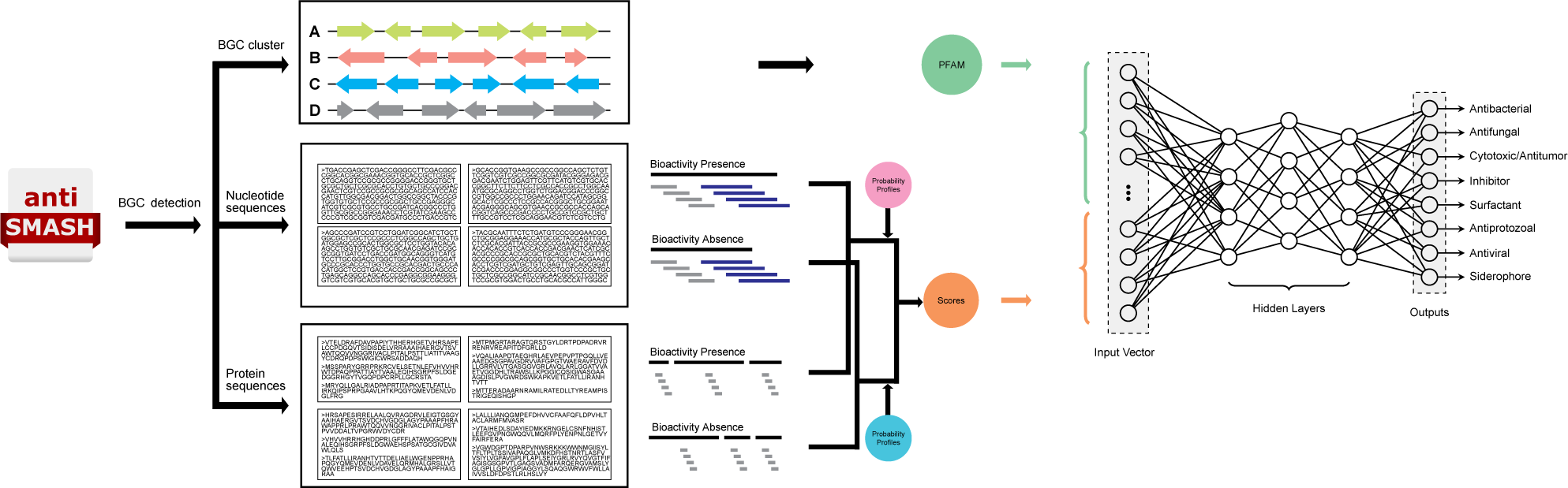
**Neural network model for bioactivity prediction of natural products**

The model is trained using a stochastic gradient descent (SGD) optimizer with a learning rate of 0.001, momentum of 0.9, and weight decay of 0.001. The network is implemented in PyTorch and trained on CPU owing to the small network size and training dataset. The training is performed using mini-batches of 64, and total epochs are set to 100.

The class imbalance is prevalent in the training dataset, as shown in Figure 1B. To deal with it, we have utilized class weighting in the loss function. This balanced heuristic approach sets weights inversely proportional to samples in the majority and minority classes. For each bioactivity, the balancing factor is calculated as follows:

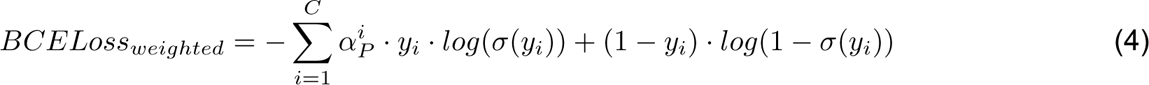

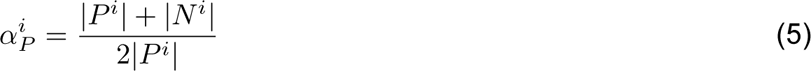

where 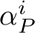 represents the loss weights for bioactivity calculated using positive (*|P ^i^|*) and negative (*|N^i^|*) instances of a class. The class weights are computed using the training set representing 90% of instances in NPB dataset.

## 3 Results

The NPBdetect tool has been developed using extensive experiments to ensure its reliability in detecting bioactivities. It includes assessing the NPF descriptors to select the best one(s) to build the model, using the latest antiSMASH tool for annotations, and integrating new sequence-based descriptors. Firstly, we aim to select a subset of descriptors from antiSMASH annotations that yield high performance. For this purpose, we performed experiments with NPF descriptors and identified PFAM as the most relevant. Subsequently, we used the latest antiSMASH tool, version 7, to extract PFAM features. Also, we have used the proposed sequence-based descriptors to extract information about the nucleotide and protein content of BGC. Both descriptors are combined, and their parameters for feature extraction are optimized to build the model. The experiments are performed using the holdout approach to develop the NPBdetect model. We have used 90% data randomly for training and the remaining for validation. The performance of the models is compared using several metrics such as accuracy, precision, recall, F1-score (FSC), and hamming loss which are given as follows:

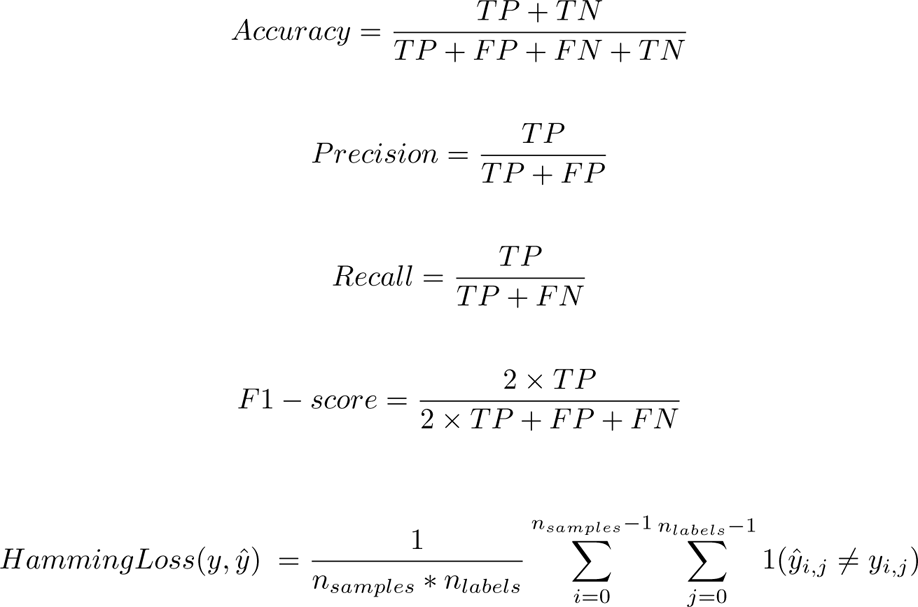

where TP denotes correctly identified bioactivity, TN denotes correctly identified bio-inactivity, FN denotes incorrectly identified bioactivity, and FP denotes incorrectly identified bio-inactivity.

### 3.1 Assessment of BGC descriptors

Although the importance of features has been determined previously when NPF tool was developed, the relevancy of individual descriptors was overlooked. Some of the descriptors may be entirely irrelevant or redundant and have a minor impact on performance for bioactivity prediction. Therefore, we have applied the feature subset selection technique on descriptors to determine relevancy. To determine the bioactivities of BGC, 14 BGC descriptors yielding 1700+ features were utilized in NPF tool. While PFAM descriptor was used in DeepBGC, the remaining ones were new to solve this problem. All descriptors except SSN and CARD can be extracted from GBKs directly. In contrast, SSNs are computed by breaking down PFAM domains into sub-PFAM domains using the SSN algorithm on the EFI-EST webserver [32, 33] and CARD features are extracted from Resistance Gene Identifier (RGI) tool [34]. Initially, we evaluated the genomic descriptors to determine their discriminatory power individually and combinatorially. For this purpose, we have used NPF tool to generate a training matrix and built ERTree and XGBoost classifiers to measure performances with 10-fold CV. The experiments are performed using antibacterial, antifungal, cytotoxic/antitumor, and antifungal/antitumor/cytotoxic classes. The ERTree classifier is used because it emerged as the best classifier in the study. In contrast, XGBoost is used as a control as it tends to generalize better than conventional ML techniques [35, 36, 37, 38].

Figure 5 presents the comparative analysis of descriptors and classifiers on the NPF dataset. Figures 5A and 5C present the outcomes of individual descriptors for ERTree and XGBoost classifiers respectively. Overall, both classifiers showed a trend for discriminating binary problems where PFAM emerged as the best, whereas NRP predicat descriptor showed the least performance. smCOG descriptor that represents the functional annotation of all accessory genes within the gene clusters also attained high accuracy but is not comparable to PFAM. Also, all variants of PK and NRP descriptors that target genes responsible for NRPS and PKS synthesis in GBKs have shown very low classification performance. The performance of non-GBK descriptors is also low as compared to PFAM. SSN descriptor attained lower accuracy than PFAM domains despite relying on them. This finding is also supported in [21] where it is shown to be redundant. Similarly, the classification accuracy of CARD descriptor is also low.

**Figure 5:**
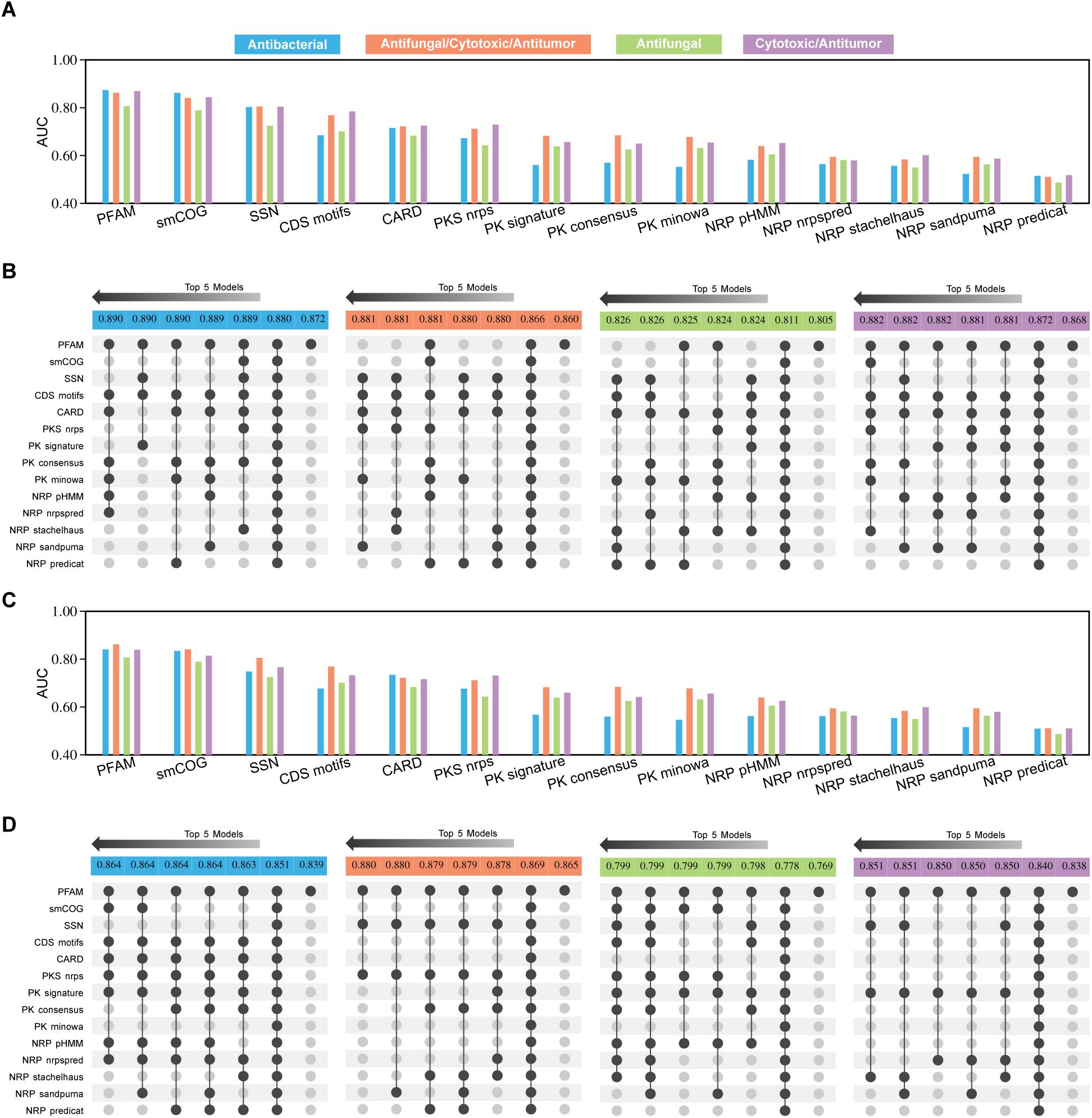
Performance analysis on NPF dataset suggesting low discriminatory power of most descriptors for bioactivity prediction. (A) Descriptor-wise performance and (B) performance of top 5 subsets of descriptors, in terms of AUC and their comparison with all descriptors as well as best descriptor (i.e., PFAM) for ERTree classifier. (C) Descriptor-wise performance and (D) performance of top 5 subsets of descriptors and their comparison with all descriptors as well as best descriptor for XGBoost classifier

Further, we determined the relevance of descriptors using a grid search approach. All combinations of descriptors were evaluated on ERTree and XGBoost classifiers, and the outcomes of the top 5 subsets are presented in Figure 5B and 5D, respectively. For comparison purposes, outcomes include all descriptors as well as the best descriptor (i.e., PFAM). The corresponding subset of descriptors is shown below the AUC outcomes. Overall, the outcomes show that several descriptors are irrelevant, and PFAM plays a vital role in attaining high performance. In all experiments with both classifiers, the best set of descriptors does not yield the best performance. Instead, descriptors are specific and contribute little to increasing performance, as observed from the top 5 subsets. Considering the best subset for all problems, all descriptors except the PK signature have been selected at least once with ERTree classifier. Likewise, all descriptors except PK minowa have been selected for the XGBoost classifier. Further, there is a minor increase in accuracy compared to PFAM for all binary problems. The maximum improvement attained with ERTree classifier is 2.16% for antifungal whereas XGBoost classifier shows an improvement of 2.57% for the antibacterial. This marginal increase shows the lack of diverse characteristics captured by other descriptors to identify the bioactivities correctly. SSN and CARD descriptors are irrelevant as subsets without them have shown similar performances for both classifiers. Similarly, other specific descriptors such as PK and NRP have a minor contribution to better decision-making for classifiers.

We have also performed experiments with NPF-filtered and NPF-latest datasets and presented the results in Supplementary Figures S1 and S2, respectively. The SSN and CARD descriptors have been excluded because of their irrelevancy. On NPF-filtered dataset (with 1191 features), the outcomes are the same as the original dataset, thereby validating the relevance of descriptors as discussed earlier. A similar trend has been observed for NPF latest dataset (1157 features), except for minor changes for antifungal datasets which could be due to fewer instances. Further, experiments are also performed to check the model overfitting due to small training dataset, and the obtained outcomes are provided in Supplementary Figure S3.

### 3.2 Neural networks for activity prediction

The neural network model to predict the bioactivities is developed by performing experiments to optimize the features. We focused on feature extraction because these handcrafted descriptors play a pivotal role in recognizing bioactivities, as shown earlier. Thus, we have fixed the model architecture and optimized the parameters of PFAM and sequence descriptors. Initially, we tested PFAM descriptor which has two parameters: similarity threshold (PFAM score) and domain frequency. We have tried score threshold from 10 to 100 using a step size of 10 and frequency threshold from 1 to 10 using a step size of 1. Figure 6 presents the outcomes for training and validation sets in terms of BCE loss. Overall, the similarity threshold is more relevant than the frequency-based threshold; however, lower similarity thresholds result in overfitting. The lower thresholds for PFAM domain selection result in high data dimensionality and vice-versa. As a result, models converged better, but validation loss remains high. The outcome is expected because of highly imbalanced classes, i.e., 5 out of 8 that affect the loss. Therefore, the selection of optimal parameters is a tedious task. The parameters that produce the higher number of features, i.e., low similarity threshold and occurrence of PFAM domains, result in low training loss but high validation loss. On the other hand, the high similarity threshold with the high occurrence of PFAM domains results in high training and validation losses.

**Figure 6:**
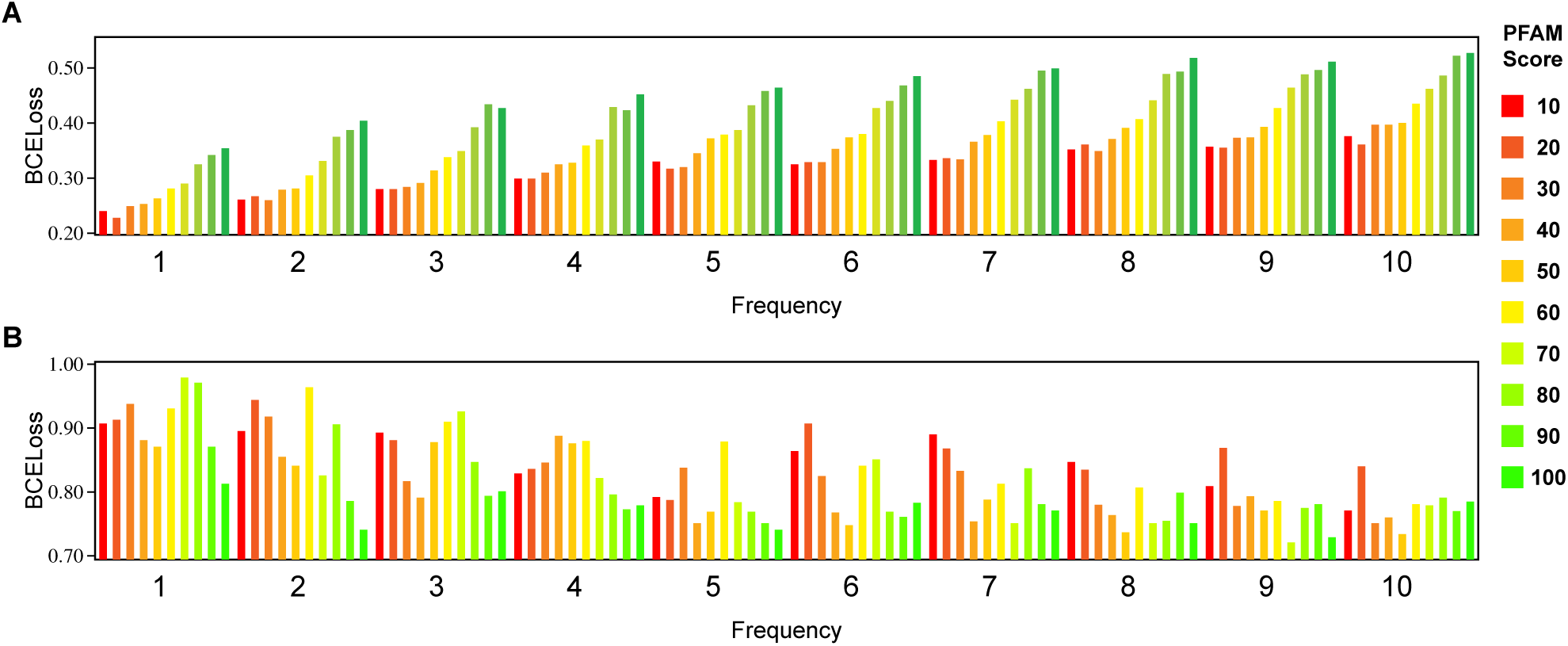
Impact of PFAM descriptor for bioactivity prediction. Features from PFAM descriptor are obtained using the similarity threshold and frequency of PFAM domains. (A) BCE loss on the training set. (B) BCE loss on the validation set.

Figure 7 presents the outcomes of *k*-mer profiles of nucleotide and peptide sequences of BGCs. For the nucleotide profiles, the training and validation loss decreases as the *k* value increases. The protein profiles also show better performance with a high *k* value while validation loss remains the same.

**Figure 7:**
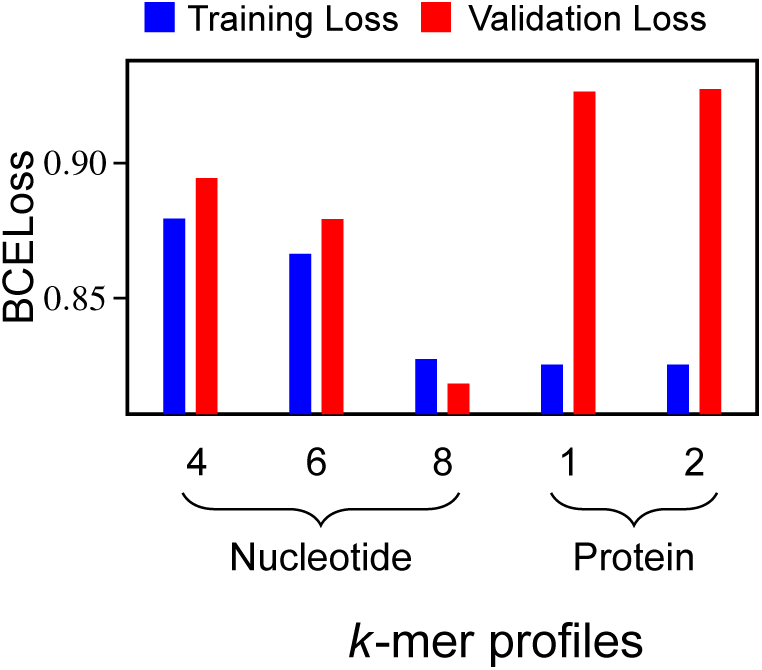
Impact of *k*-mer profiles for bioactivity prediction. For nucleotide sequence of BGCs, 4, 6 & 8-mers are computed. For protein sequences of BGCs, 1 & 2-mers are computed.

Taking motivation from feature subset selection [39] that high relevancy does not necessarily imply that a descriptor is optimal, PFAM and sequence composition descriptors are combined to optimize the parameters. We have excluded the high-frequency parameter (>5) and used the low threshold for PFAM domain selection (<50) for further experiments as a higher value results in the loss of essential information that makes the classifier more sensitive to the detection of similar BGCs and thereby could miss the novel ones.

We have performed experiments with various combinations of PFAM and sequence-based descriptors, including nucleotide profiles (NP) and protein profiles (PP). In total, seven combinations are tried which include combination of PFAM with (1) nucleotide profiles only, (2) all peptide profiles only, (3) all nucleotide as well as peptide profiles, (4) 6 & 8-mer nucleotide as well as all peptide profiles, (5) 8-mer nucleotide as well as all peptide profiles, (6) 6 & 8-mer nucleotide as well as 2-mer peptide profiles and (7) all nucleotide profiles as well as 2-mer peptide profiles. The best parametric value is determined using the lower validation losses and their difference between corresponding training losses. Overall, a trend has been found where training loss is higher while validation loss becomes lesser when PFAM domain cutoff score is 40, as shown in Figure 8 as well as other combinations provided in Supplementary Figures S4-S9. Even though the differences are minor for the best PFAM domain cutoff, selecting frequency is tedious as combinations with nucleotide and protein profiles have yielded similar outcomes. So, we have selected the 6 & 8-mer nucleotide and 2-mer peptide profiles among all these sequence descriptor combinations because of slightly better performance.

**Figure 8:**
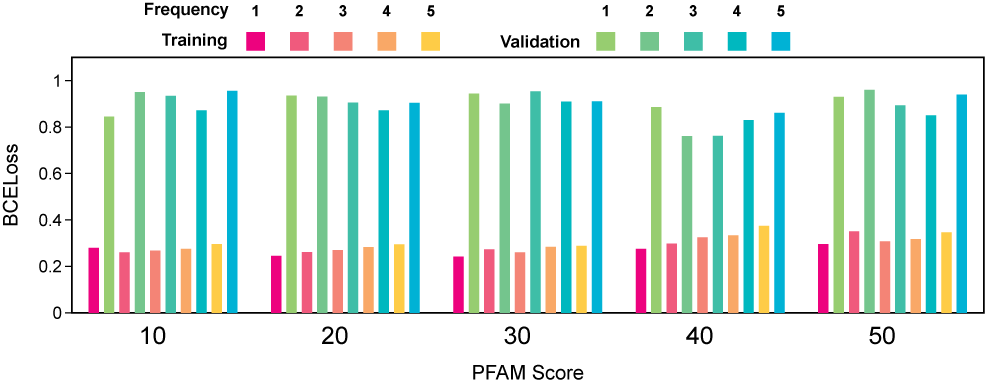
Selection of PFAM threshold cutoff using the best combination of sequence descriptors (nucleotide profiles with 6 and 8 mer and protein profile with 2 mer)

Figure 8 presents the outcomes of the combination of PFAM and sequence descriptors. At the lower PFAM frequency thresholds (1, 2, and 3) with all scores, overfitting is common as training loss is less while validation loss is high compared to the higher frequency cutoffs and scores. At the frequency threshold of 4, the training and validation losses are consistent while training and validation losses for frequency threshold 5 are comparatively higher. Thus, using the minimum difference between training and validation loss as a criterion for selection, we selected a score threshold of 40 and a frequency cutoff of 4 to be optimal.

### 3.3 Bioactivity prediction on NPB-LM test set

To validate the efficacy of NPBdetect tool, we have tested the model on the NPB-LM test set. The obtained outcomes are presented in Figure 9 in terms of performance metrics, AUC plots, and confusion matrices for all bioactivities. Overall, we have observed that NPBdetect tool showed reasonable performance on the test set for most bioactivities. Figure 9A presents the outcomes for all bioactivities in terms of balanced accuracy, precision, recall, F1-score, and AUC values. Despite the small dataset size, the model identified siderophore bioactivity with the highest accuracy, whereas antibacterial, antifungal, and cytotoxic/antitumor bioactivities are recognized with reasonable accuracy. The most challenging bioactivities are surfactant, antiprotozoal, and antiviral, whose instances remain undetected. Except for antibacterial and siderophore bioactivities, the precision of the remaining bioactivities is very low, which indicates that the model tends to have a high false positive rate. In terms of recall rate, the model detected cytotoxic/antitumor and siderophore bioactivities correctly while identification of antibacterial, antifungal and inhibitor bioactivities remained low and the rest of the classes are not detected at all. A similar trend has been observed for FSC where high score is obtained for cytotoxic/antitumor and siderophore bioactivities. The identifications for the rest of the bioactivities, namely inhibitor, surfactant, antiprotozoal, and antiviral, are incorrect. The poor performance can be attributed to high class-imbalance as well as the absence of quality patterns in extracted features to detect these bioactivities.

**Figure 9:**
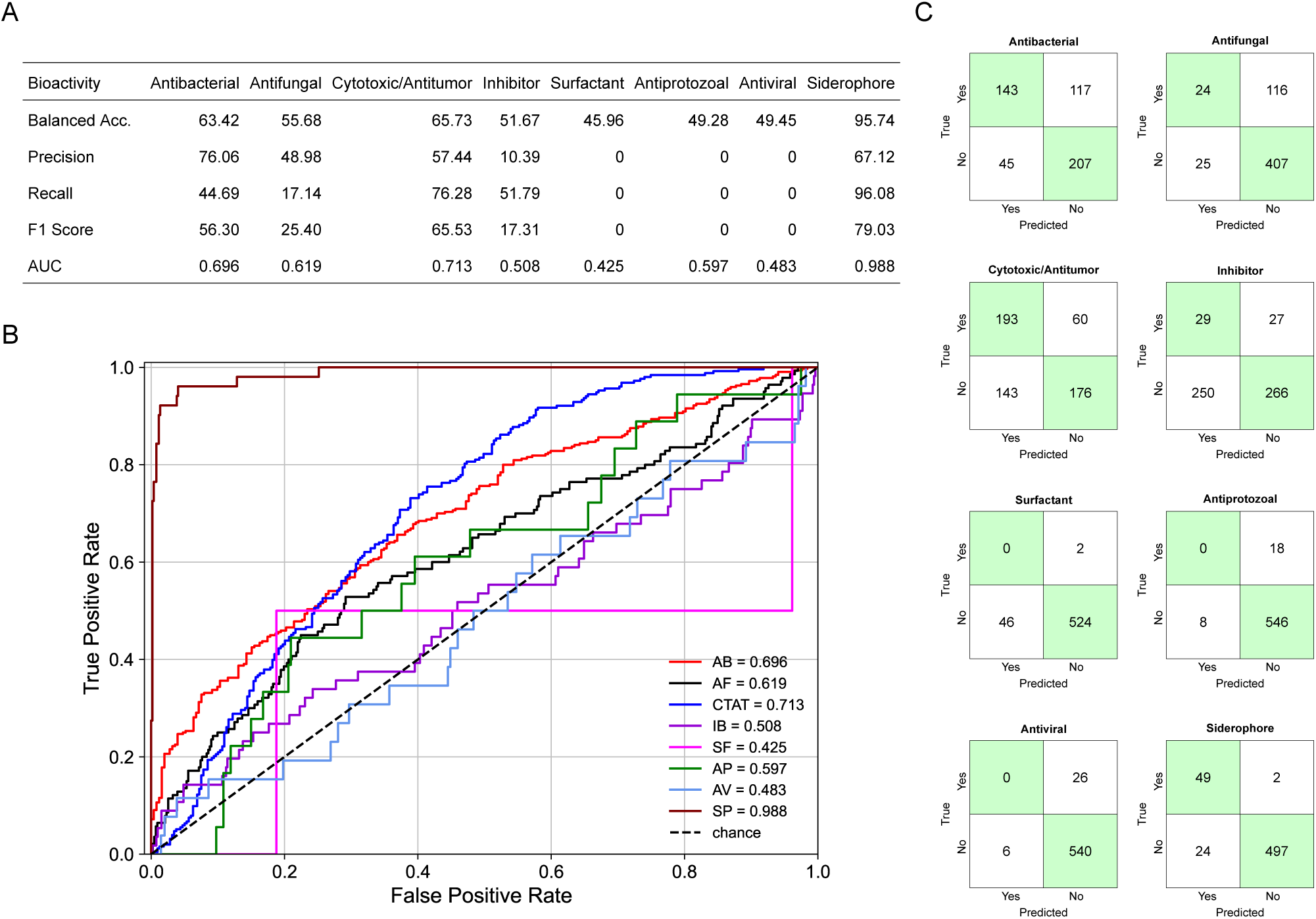
Performance of NPBdetect tool on NPF-LM test set. (A) Performance in terms of balanced accuracy, precision, recall, F1-score, and AUC. (B) ROC curve for all bioactivities. (C) Confusion matrices for all bioactivities.

Figure 9B presents the ROC curves for all bioactivities. The key finding from these outcomes is that predictions for inhibitor, antiprotozoal, antiviral, and surfactant bioactivities are similar to a random classifier. Figure 9C shows the confusion matrix for each bioactivity. The model is inclined towards the majority class, i.e., lack of bioactivity for the highly-imbalanced classes such as surfactant, antiprotozoal, and antiviral. As a result, no instances of these bioactivities are detected. Siderophore is an exception in high-imbalanced classes whose majority of instances are recognized correctly. Antibacteria and antifungal bioactivities also showed the same trend where most predictions are of the absence of bioactivity. On the other hand, the model is inclined toward the presence of bioactivity for cytotoxic/antitumor bioactivity and, thus, has a high detection rate. In the case of inhibitor, the detection rate for both the presence and absence of bioactivity is equal which may signify the random predictions by the model. The hamming loss for all bioactivities is 0.20 which indicates room for improvement as misclassifications are high.

### 3.4 Comparison with previous work

The NPBdetect tool is also compared with the NPF tool on NPB-LM dataset to validate its better generalization and superiority. The comparison is made using antibacterial, antifungal and cytotoxic/antitumor bioactivities. The antigram-positive, antigram-negative, and cytotoxic/antitumor/antifungal are excluded because these are subsets of antibacterial, antifungal, and cytotoxic classes. Also, comparisons are made using two versions of antiSMASH and different classifiers used in NPF tool. The obtained outcomes are presented in Figure 10 in terms of balanced accuracy, precision, recall, F1-score, and AUC. Initially, NPB-LM test dataset is evaluated by processing BGCs through antiSMASH 4. The model is trained using NPF-filtered instances with all descriptors except SSN. The outcomes show the lack of the best classifier among ERTree, SVM with RBF kernel, and SGD classifiers, as shown in Figure 10A. The SVM-RBF classifier attained the best-balanced accuracy and F1-score for antibacterial and cytotoxic/antitumor bioactivities whereas SGD classifier achieved the best AUC for all three bioactivities. ERTree classifier outperformed the other two classifiers for antifungal bioactivity. Also, SVM-RBF classifier has the best whereas SGD classifier has the lowest recall rate. Overall, we concluded that no classifier is better than others, which validates the need to develop better models. Also, performance bottlenecks are observed due to lack of sufficient data.

**Figure 10:**
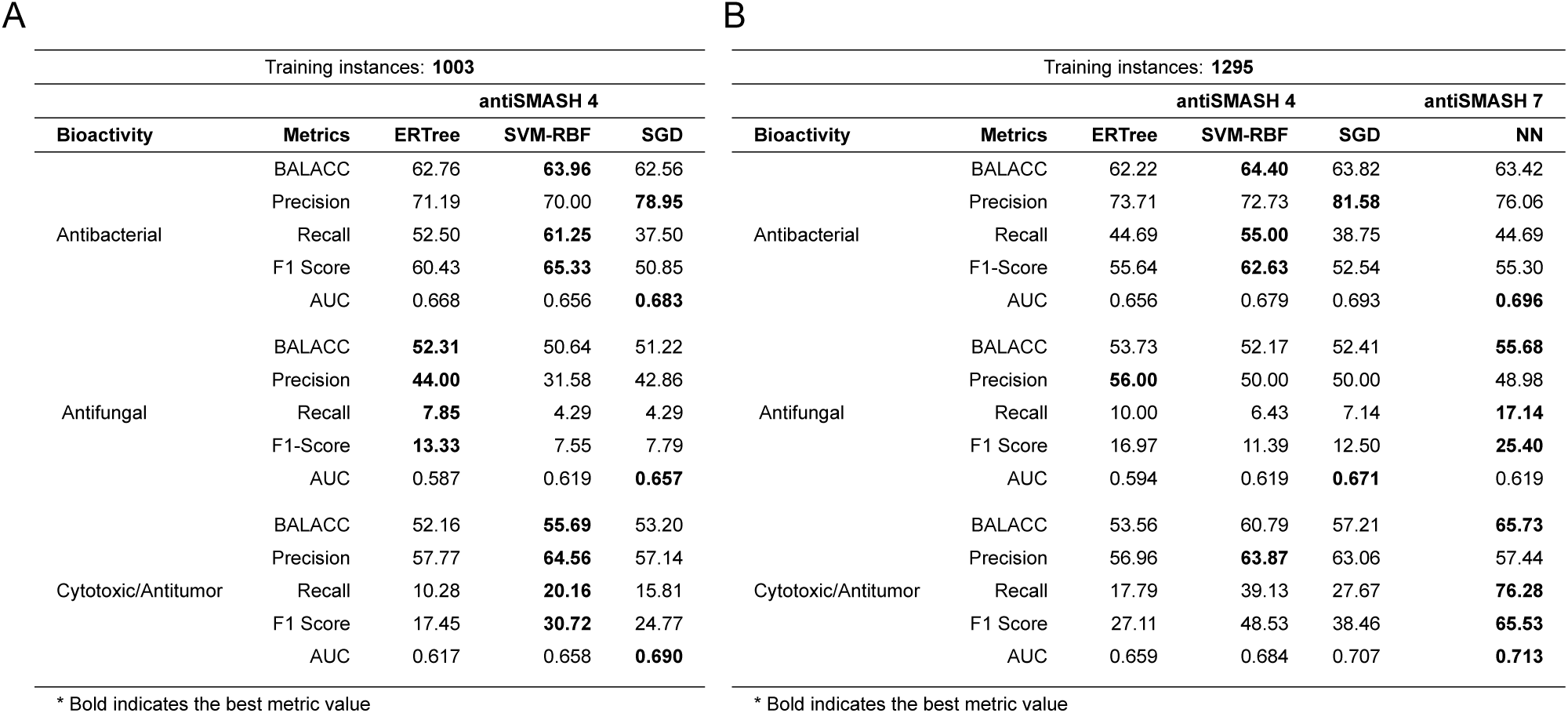
Comparison of NPBdetect and NPF tools. (A) Comparison of NPF classifiers on NPB-LM test set. (B) Comparison of NPF and NPBdetect classifiers trained on NPB training set and model validation on NPB-LM test set.

Further, we extended NPF models using the latest training dataset of 1295 instances. The new training dataset is also processed through antiSMASH 4, and all descriptors except SSN were used to build models with the same three classifiers using pre-defined parameters. On the other hand, NPBdetect tool used the antiSMASH 7 processed BGCs. The classifiers are evaluated on NPB-LM test set, and their outcomes are presented in Figure 10B. Overall, NPBdetect tool outperformed the NPF tool, and a larger training set also improved the model accuracy of antiSMASH 4 processed BGCs. The proposed combination of PFAM domains as well as BGC nucleotide and protein sequences information help to build a superior model. The NPF tool was outperformed with an improved balanced accuracy of approx. 2%, and 5% for antifungal and cytotoxic/antitumor bioactivities respectively. On the other hand, the outcomes of antibacterial bioactivity are comparable with best of three NPF classifiers. The differences in the precision of the best NPF classifier and NPBdetect tool are approx. 5%, 8%, and 6% for antibacterial, antifungal and cytotoxic/antitumor bioactivities respectively. However, the recall rate of NPBdetect is significantly higher than the best of three classifiers, i.e., 7% and 37% for antifungal and cytotoxic/antitumor bioactivities respectively. Similarly, NPBdetect tool also showed the best F1-score with an improvement of 8% and 17% for antifungal and cytotoxic/antitumor bioactivities respectively. In terms of AUC, it showed higher AUC on antibacterial and cytotoxic/antitumor bioactivities whereas SGD classifier has shown better AUC for antifungal activity, but with a minor improvement of 0.05.

Comparing NPF classifiers only, the outcomes are similar to previous findings, i.e., no classifier is better than others. These classifiers have shown higher precision than the NPBdetect, but the differences are comparable. Additionally, higher quality and larger training set, after removing instances as per the latest MIBiG database and adding new BGC instances, also improved the performance of NPF tool on three classifiers. This finding validates the need to increase the size of training datasets to build better models that can be utilized to search bioactivities at a large scale. Thus, it can be concluded that the NPBdetect tool outperformed the NPF tool with three sub-optimal classifiers.

### 3.5 Evaluating false positives and new compounds

To check the reliability of the model in capturing patterns for detecting bioactivities, it is essential to test the model predictions through experiments. This activity establishes the model’s confidence to discover bioactivities correctly and rule out the possibility of random or biased predictions by models, as noted for four classes earlier. To achieve this, we selected false positives and turned to the latest literature again to find bioactivities validated by researchers in recent experiments. The false positives are selected because the lack of bioactivity in NPB-LM indicates either the absence of activity or the lack of experimental validation. Specifically, false positives of antibacterial, antifungal, and cytotoxic/antitumor identified by NPBdetect were searched. Interestingly, we got multiple hits where misclassified BGCs were indeed bioactive. It includes 2 antibacterial [40, 41], 1 antifungal [42] and 4 cytotoxic/antitumor [43, 44, 45, 46] predictions. Moreover, we found that the predictions of these instances are similar to NPF tool (i.e., false positives) which also substantiates the reliability of the proposed tool.

Further, we checked the siderophore bioactivity as it has been detected with the highest accuracy by the model. Through literature, we found three new siderophores with compounds, namely *Desferriferrichrome* [47], *Fradiamines* [48], and *Rhizoferrin* [49]. These compounds are not part of either NPB training or test sets and are validated by researchers through in-vitro experiments. The sequences are processed through antiSMASH, and obtained BGCs are evaluated on NPBdetect tool which classified 2 out of 3 compounds as siderophore. These outcomes validate the model’s reliability in identifying the four bioactivities with high confidence and usability of the tool for real-world applications to detect the bioactivities from BGCs. Moreover, we used the model to discover the bioactivites of 635 unannotated BGCs of MIBiG database which do not overlap with either training or test sets. The predictions are provided in Supplementary Table S2, and their respective class distribution is in Supplementary Figure S10.

## 4 Discussion

Natural products have emerged as an essential research domain to tackle the increasing antibiotic resistance across the globe [50, 51]. Thus, their proven clinical and therapeutic applications require accelerated identification of bioactivities, and AI-driven computational approach has the potential to streamline and expedite the identification at a large scale. These approaches can reduce the experimental time and cost substantially to confirm the bioactivities through experiments. However, we found that predicting the functions of natural products is a challenging task, from data collection to model building.

Firstly, the lack of integrated and curated databases and non-standard naming conventions [52] is the most prominent challenge that could result in erroneous labels during data curation. For instance, we found that many studies had mentioned antibiotics or antimicrobials for both antibacterial and antifungal activities. We resolved this issue by looking for organisms on which bioactivity was tested and then decided on the label as antibacterial or antifungal. The standardization of the naming of bioactivities by using the MIBiG database as a reference could make the curation process easier in future research [53]. Also, more experimental validation is required to capture the bioactivities and incorporate the resultant information in ML datasets.

Secondly, antiSMASH tool has been employed to annotate BGCs in this work. This tool is developed with the incremental approach where new releases add new rules and update old ones, affecting the feature extraction from descriptors present in GBK files. For instance, NRPS and PKS annotations are found in *sec_met* tag of antiSMASH 4, but this information is located in *aSDomain* tag of antiSMASH 7. Also, some new tags are introduced in antiSMASH 7, including *gene_ontologies*, *gene_functions*, *SMILES*, and *monomer_pairings*. This incremental build approach demands keeping up with the latest computational methods to mine BGCs from genome and metagenome sequences and re-do the feature extraction and model training, including parameter optimization of descriptors as well as refinements of network architecture.

Lastly, the low detection rate of inhibitor, surfactant, antiprotozoal, and antiviral bioactivities shows the limitation of this work. These bioactivities comprise half of the classes tried in this work. The surfactant, antivi-ral, and antiprotozoal bioactivities are the toughest, whereas inhibitor bioactivity may be detected by chance only. While these outcomes may be attributed to class-imbalance, the model recognized the similar bioactivity, siderophore, with high accuracy. We believe this happened as the proposed descriptors captured the patterns well for siderophore while failing to do so for the remaining classes. This opens up the possibility of designing more descriptors to improve the performance of models developed with ML techniques. In addition, deep learning models have not yet been explored for this multi-label classification problem.

## 5 Conclusion

We developed a neural network model called NPBdetect to detect the bioactivities of natural products from their BGCs. This multi-label classification problem, which has gained the focus of ML community recently, is challenging due to the number of shortcomings like small datasets, lack of relevant descriptors, and capability to detect few bioactivities only. In this work, we have taken several steps to address this problem effectively which includes creating a large training set by refining the older one and using the latest available information, creating a test set using literature curation, evaluating existing descriptors to select the best one and proposing new sequence based descriptor and extending the model to detect a higher number of bioactivities than previous attempts. The evaluation of the independent test set showed that NPBdetect model performed well overall by attaining high accuracy for a highly imbalanced class, reasonable accuracy for three classes, and low accuracy for the remaining four highly imbalanced classes. Also, it outperformed the NPF tool in several experiments which shows its effectiveness in detecting bioactivities better.

This work also opens up opportunities for the biology community to make efforts for data curation to build larger datasets and for the ML community to come up with more relevant descriptors and develop strategies to deal with class-imbalance.

## 6 Data Availability

The code and data is available on request to authors.

## 7 Acknowledgements

We would like to thank National Agri-Food Biotechnology Institute (NABI) for providing us with infrastructure and supporting high-performance computing. We acknowledge National Supercomputing Mission (NSM) for providing computing resources of **PARAM Smriti** at NABI, Mohali, which is implemented by C-DAC and supported by the Ministry of Electronics and Information Technology (MeitY), Department of Biotechnology (DBT) and Department of Science and Technology (DST), Government of India. We also acknowledge DeLCON (DBTelectronic library consortium), Gurugram, India, for the online journal access.

Shrikant Mantri would like to thank Professor Nadine Ziemert for insightful discussion about this topic during the conception stage of this study and acknowledges NABI Core funding to execute this project. Hemant Goyat acknowledge NABI for his project assistant position. Sunaina Paliyal acknowledge NABI for her computational biology training extension to continue working on this project.

## Author contributions

H.G., D.S. and S.M. conceptualised and designed the study. H.G. and S.P. conducted the literature review and collected data. D.S. developed the model. D.S. and H.G. conducted the experiments and analysed the results. H.G., D.S. and S.P. wrote the manuscript. S.M. supervised and reviewed the manuscript.

## References

[1] Marnix H Medema, Renzo Kottmann, Pelin Yilmaz, Matthew Cummings, John B Biggins, Kai Blin, Irene De Bruijn, Yit Heng Chooi, Jan Claesen, R Cameron Coates, et al. Minimum information about a biosynthetic gene cluster. Nature chemical biology, 11(9):625–631, 2015.

[2] Arnold L Demain and Aiqi Fang. The natural functions of secondary metabolites. History of modern biotechnology I, pages 1–39, 2000.

[3] Ibrahim Ilker Ozyigit, Ilhan Dogan, Asli Hocaoglu-Ozyigit, Bestenur Yalcin, Aysegul Erdogan, Ibrahim Ertugrul Yalcin, Evren Cabi, and Yilmaz Kaya. Production of secondary metabolites using tissue culture-based biotechnological applications. Frontiers in Plant Science, 14:1132555, 2023.

[4] R Dieckmann, I Graeber, I Kaesler, U Szewzyk, and H Von Döhren. Rapid screening and dereplication of bacterial isolates from marine sponges of the sula ridge by intact-cell-maldi-tof mass spectrometry (icm-ms). Applied microbiology and biotechnology, 67:539–548, 2005.

[5] Tristan Richard, Hamza Temsamani, Emma Cantos-Villar, and Jean-Pierre Monti. Application of lc–ms and lc–nmr techniques for secondary metabolite identification. In Advances in botanical research, volume 67, pages 67–98. Elsevier, 2013.

[6] Christopher G Jones, Michael W Martynowycz, Johan Hattne, Tyler J Fulton, Brian M Stoltz, Jose A Rodriguez, Hosea M Nelson, and Tamir Gonen. The cryoem method microed as a powerful tool for small molecule structure determination. ACS Central Science, 4(11):1587–1592, 2018.

[7] Lee Joon Kim, Masao Ohashi, Zhuan Zhang, Dan Tan, Matthew Asay, Duilio Cascio, José A Rodriguez, Yi Tang, and Hosea M Nelson. Prospecting for natural products by genome mining and microcrystal electron diffraction. Nature chemical biology, 17(8):872–877, 2021.

[8] Satria A Kautsar, Kai Blin, Simon Shaw, Tilmann Weber, and Marnix H Medema. Big-fam: the biosynthetic gene cluster families database. Nucleic acids research, 49(D1):D490–D497, 2021.

[9] Michael W Mullowney, Katherine R Duncan, Somayah S Elsayed, Neha Garg, Justin JJ van der Hooft, Nathaniel I Martin, David Meijer, Barbara R Terlouw, Friederike Biermann, Kai Blin, et al. Artificial intelligence for natural product drug discovery. Nature Reviews Drug Discovery, 22(11):895–916, 2023.

[10] Katherine D Bauman, Keelie S Butler, Bradley S Moore, and Jonathan R Chekan. Genome mining methods to discover bioactive natural products. Natural Product Reports, 38(11):2100–2129, 2021.

[11] Kai Blin, Simon Shaw, Katharina Steinke, Rasmus Villebro, Nadine Ziemert, Sang Yup Lee, Marnix H Medema, and Tilmann Weber. antismash 5.0: updates to the secondary metabolite genome mining pipeline. Nucleic acids research, 47(W1):W81–W87, 2019.

[12] Geoffrey D Hannigan, David Prihoda, Andrej Palicka, Jindrich Soukup, Ondrej Klempir, Lena Rampula, Jindrich Durcak, Michael Wurst, Jakub Kotowski, Dan Chang, et al. A deep learning genome-mining strategy for biosynthetic gene cluster prediction. Nucleic acids research, 47(18):e110–e110, 2019.

[13] Michael A Skinnider, Nishanth J Merwin, Chad W Johnston, and Nathan A Magarvey. Prism 3: expanded prediction of natural product chemical structures from microbial genomes. Nucleic acids research, 45(W1):W49–W54, 2017.

[14] Jinki Kim and Gwan-Su Yi. Pkminer: a database for exploring type ii polyketide synthases. BMC microbiology, 12(1):1–12, 2012.

[15] Priyesh Agrawal, Shradha Khater, Money Gupta, Neetu Sain, and Debasisa Mohanty. Rippminer: a bioinformatics resource for deciphering chemical structures of ripps based on prediction of cleavage and cross-links. Nucleic acids research, 45(W1):W80–W88, 2017.

[16] Jonathan I Tietz, Christopher J Schwalen, Parth S Patel, Tucker Maxson, Patricia M Blair, Hua-Chia Tai, Uzma I Zakai, and Douglas A Mitchell. A new genome-mining tool redefines the lasso peptide biosynthetic landscape. Nature chemical biology, 13(5):470–478, 2017.

[17] Emmanuel LC de Los Santos. Neuripp: Neural network identification of ripp precursor peptides. Scientific reports, 9(1):13406, 2019.

[18] Nishanth J Merwin, Walaa K Mousa, Chris A Dejong, Michael A Skinnider, Michael J Cannon, Haoxin Li, Keshav Dial, Mathusan Gunabalasingam, Chad Johnston, and Nathan A Magarvey. Deepripp integrates multiomics data to automate discovery of novel ribosomally synthesized natural products. Proceedings of the National Academy of Sciences, 117(1):371–380, 2020.

[19] Auke J van Heel, Anne de Jong, Chunxu Song, Jakob H Viel, Jan Kok, and Oscar P Kuipers. Bagel4: a user-friendly web server to thoroughly mine ripps and bacteriocins. Nucleic acids research, 46(W1):W278– W281, 2018.

[20] Allison S Walker and Jon Clardy. A machine learning bioinformatics method to predict biological activity from biosynthetic gene clusters. Journal of Chemical Information and Modeling, 61(6):2560–2571, 2021.

[21] Olivia Riedling, Allison S Walker, and Antonis Rokas. Predicting fungal secondary metabolite activity from biosynthetic gene cluster data using machine learning. Microbiology Spectrum, 12(2):e03400–23, 2024.

[22] Ruihan Zhang, Xiaoli Li, Xingjie Zhang, Huayan Qin, and Weilie Xiao. Machine learning approaches for elucidating the biological effects of natural products. Natural Product Reports, 38(2):346–361, 2021.

[23] Barbara R Terlouw, Kai Blin, Jorge C Navarro-Munoz, Nicole E Avalon, Marc G Chevrette, Susan Egbert, Sanghoon Lee, David Meijer, Michael JJ Recchia, Zachary L Reitz, et al. Mibig 3.0: a community-driven effort to annotate experimentally validated biosynthetic gene clusters. Nucleic acids research, 51(D1):D603– D610, 2023.

[24] Kai Blin, Simon Shaw, Hannah E Augustijn, Zachary L Reitz, Friederike Biermann, Mohammad Alanjary, Artem Fetter, Barbara R Terlouw, William W Metcalf, Eric JN Helfrich, et al. antismash 7.0: new and improved predictions for detection, regulation, chemical structures and visualisation. Nucleic acids research, 51(W1):W46–W50, 2023.

[25] Liguo Wang, Hyun Jung Park, Surendra Dasari, Shengqin Wang, Jean-Pierre Kocher, and Wei Li. Cpat: Coding-potential assessment tool using an alignment-free logistic regression model. Nucleic acids research, 41(6):e74–e74, 2013.

[26] Valentin Wucher, Fabrice Legeai, Benoît Hédan, Guillaume Rizk, Lætitia Lagoutte, Tosso Leeb, Vidhya Jagannathan, Edouard Cadieu, Audrey David, Hannes Lohi, et al. Feelnc: a tool for long non-coding rna annotation and its application to the dog transcriptome. Nucleic acids research, 45(8):e57–e57, 2017.

[27] Dalwinder Singh and Joy Roy. A large-scale benchmark study of tools for the classification of protein-coding and non-coding rnas. Nucleic Acids Research, 50(21):12094–12111, 2022.

[28] Ian Goodfellow, Yoshua Bengio, and Aaron Courville. Deep learning. MIT press, 2016.

[29] Josh Patterson and Adam Gibson. Deep learning: A practitioner’s approach.” O’Reilly Media, Inc.”, 2017.

[30] Nishant Ravikumar, Arezoo Zakeri, Yan Xia, and Alejandro F Frangi. Deep learning fundamentals. In Medical Image Analysis, pages 415–450. Elsevier, 2024.

[31] Dan Hendrycks and Kevin Gimpel. Gaussian error linear units (gelus). arXiv preprint arXiv:1606.08415, 2016.

[32] John A Gerlt, Jason T Bouvier, Daniel B Davidson, Heidi J Imker, Boris Sadkhin, David R Slater, and Katie L Whalen. Enzyme function initiative-enzyme similarity tool (efi-est): a web tool for generating protein sequence similarity networks. Biochimica Et Biophysica Acta (BBA)-Proteins and Proteomics, 1854(8):1019–1037, 2015.

[33] Nils Oberg, Rémi Zallot, and John A Gerlt. Efi-est, efi-gnt, and efi-cgfp: enzyme function initiative (efi) web resource for genomic enzymology tools. Journal of molecular biology, 435(14):168018, 2023.

[34] Brian P Alcock, Amogelang R Raphenya, Tammy TY Lau, Kara K Tsang, Mégane Bouchard, Arman Edalatmand, William Huynh, Anna-Lisa V Nguyen, Annie A Cheng, Sihan Liu, et al. Card 2020: antibiotic resistome surveillance with the comprehensive antibiotic resistance database. Nucleic acids research, 48(D1):D517–D525, 2020.

[35] Yijun Zhao, Tong Wang, Riley Bove, Bruce Cree, Roland Henry, Hrishikesh Lokhande, Mariann Polgar-Turcsanyi, Mark Anderson, Rohit Bakshi, Howard L Weiner, et al. Ensemble learning predicts multiple sclerosis disease course in the summit study. NPJ digital medicine, 3(1):135, 2020.

[36] Adeola Ogunleye and Qing-Guo Wang. Xgboost model for chronic kidney disease diagnosis. IEEE/ACM transactions on computational biology and bioinformatics, 17(6):2131–2140, 2019.

[37] Ravid Shwartz-Ziv and Amitai Armon. Tabular data: Deep learning is not all you need. Information Fusion, 81:84–90, 2022.

[38] Léo Grinsztajn, Edouard Oyallon, and Gaël Varoquaux. Why do tree-based models still outperform deep learning on typical tabular data? Advances in Neural Information Processing Systems, 35:507–520, 2022.

[39] Ron Kohavi and George H John. Wrappers for feature subset selection. Artificial intelligence, 97(1-2):273– 324, 1997.

[40] Ryuichi Sawa, Yoshiaki Takahashi, Hideki Hashizume, Kazushige Sasaki, Yoshimasa Ishizaki, Maya Umekita, Masaki Hatano, Hikaru Abe, Takumi Watanabe, Naoko Kinoshita, et al. Amycolamicin: a novel broad-spectrum antibiotic inhibiting bacterial topoisomerase. Chemistry–A European Journal, 18(49):15772–15781, 2012.

[41] Yasushi Ogasawara, Shuhei Umetsu, Yuki Inahashi, Kenichi Nonaka, and Tohru Dairi. Identification of pulvomycin as an inhibitor of the futalosine pathway. The Journal of Antibiotics, 74(11):825–829, 2021.

[42] Rupanshee Srivastava, Rajesh Prajapati, Tripti Kanda, Sadhana Yadav, Nidhi Singh, Shivam Yadav, Rajeev Mishra, and Neelam Atri. Phycochemistry and bioactivity of cyanobacterial secondary metabolites. Molecular biology reports, 49(11):11149–11167, 2022.

[43] George E Chlipala, Megan Sturdy, Aleksej Krunic, Daniel D Lantvit, Qi Shen, Kyle Porter, Steven M Swanson, and Jimmy Orjala. Cylindrocyclophanes with proteasome inhibitory activity from the cyanobacterium nostoc sp. Journal of natural products, 73(9):1529–1537, 2010.

[44] Katherine Walton and John P Berry. Indole alkaloids of the stigonematales (cyanophyta): Chemical diversity, biosynthesis and biological activity. Marine Drugs, 14(4):73, 2016.

[45] Hee-Ju Nah, Jihee Park, Sisun Choi, and Eung-Soo Kim. Wbla, a global regulator of antibiotic biosynthesis in streptomyces. Journal of Industrial Microbiology and Biotechnology, 48(3-4):kuab007, 2021.

[46] Valentinos Kounnis, Georgios Chondrogiannis, Michalis D Mantzaris, Andreas G Tzakos, Demosthenes Fokas, Nikolaos A Papanikolaou, Vasiliki Galani, Ioannis Sainis, and Evangelos Briasoulis. Microcystin lr shows cytotoxic activity against pancreatic cancer cells expressing the membrane oatp1b1 and oatp1b3 transporters. Anticancer research, 35(11):5857–5865, 2015.

[47] Huiwen Liu, Liangyin Sun, Jintao Zhang, Yongzhong Wang, and Hengqian Lu. Siderophore-synthesizing nrps reprogram lipid metabolic profiles for phenotype and function changes of arthrobotrys oligospora. World Journal of Microbiology and Biotechnology, 40(2):46, 2024.

[48] Jianwei Chen, Yuqi Guo, Qihao Wu, Wei Wang, Jiangwei Pan, Minghong Chen, Hong Jiang, Qunjian Yin, Gaiyun Zhang, Bin Wei, et al. Discovery of new siderophores from a marine streptomycetes sp. via combined metabolomics and analysis of iron-chelating activity. Journal of Agricultural and Food Chemistry, 71(17):6584–6593, 2023.

[49] Alberto E Lopez, Lubov S Grigoryeva, Armando Barajas, and Nicholas P Cianciotto. Legionella pneumophila rhizoferrin promotes bacterial biofilm formation and growth within amoebae and macrophages. Infection and immunity, 91(8):e00072–23, 2023.

[50] Sean E Rossiter, Madison H Fletcher, and William M Wuest. Natural products as platforms to overcome antibiotic resistance. Chemical reviews, 117(19):12415–12474, 2017.

[51] David J Newman and Gordon M Cragg. Natural products as sources of new drugs from 1981 to 2014. Journal of natural products, 79(3):629–661, 2016.

[52] Nadja B Cech, Marnix H Medema, and Jon Clardy. Benefiting from big data in natural products: importance of preserving foundational skills and prioritizing data quality. Natural Product Reports, 38(11):1947–1953, 2021.

[53] Rajinder Gupta and Shrikant S Mantri. Biomolecular relationships discovered from biological labyrinth and lost in ocean of literature: Community efforts can rescue until automated artificial intelligence takes over. Frontiers in Genetics, 7:186589, 2016.

