## Supplementary File for "Predicting biological activity from biosynthetic gene clusters using neural networks"

#### List of Figures

S1 Performance analysis on NPP-filtered instances using ERTree and XGBoost classifiers. (A) Descriptor-wise performance and (B) performance of top 5 subsets of descriptors, in terms of AUC and their comparison with all descriptors as well as best descriptor (i.e., PFAM) for ERTree classifier. (C) Descriptor-wise performance and (D) performance of top 5 subsets of descriptors and their comparison with all descriptors as well as best descriptor for XGBoost classifier . . . . 2

S2 Performance analysis on NPP-latest instances using ExtraTree and XGBoost classifiers. (A) Descriptor-wise performance and (B) Performance of top 5 subsets of descriptors, in terms of AUC and their comparison with all descriptors as well as best descriptor (i.e., PFAM) for ERTree classifier. (C) Descriptor-wise performance and (D) performance of top 5 subsets of descriptors and their comparison with all descriptors as well as best descriptor for XGBoost classifier . . . . 3

S3 AUC outcomes for different bioactivity classes on NPF training (blue) and MIBiG test (red) sets. AB denotes antibacterial, AF denotes antifungal, and CT/AT denotes antitumor or cytotoxic . . 4

S4 Performance with combination of PFAM and sequence descriptors (nucleotide profiles with 4, 6 and 8-mers) . . . . . . . . . . . . . . . . . . . . . . . . . . . . . . . . . . . . . . . . . 5

S5 Performance with combination of PFAM and sequence descriptors (peptide profiles with 1 and 2-mers) . . . . . . . . . . . . . . . . . . . . . . . . . . . . . . . . . . . . . . . . . 6

S6 Performance with combination of PFAM and sequence descriptors (nucleotide profiles with 4, 6 and 8-mers and peptide profiles with 1 and 2-mers) . . . . . . . . . . . . . . . . . . . . . . 6

S7 Performance with combination of PFAM and sequence descriptors (nucleotide profiles with 6 and 8-mers and peptide profiles with 1 and 2-mers) . . . . . . . . . . . . . . . . . . . . . . 6

S8 Performance with combination of PFAM and sequence descriptors (nucleotide profile with 8 mer and peptide profiles with 1 and 2-mers) . . . . . . . . . . . . . . . . . . . . . . . . . 7

S9 Performance with combination of PFAM and sequence descriptors (nucleotide profile with 4, 6 and 8-mers and peptide profiles with 2-mers) . . . . . . . . . . . . . . . . . . . . . . . . . 7

S10 Distribution of the predicted Bioactivities for 635 unannotated MIBiG BGCs . . . . . . . . . . . 8

### 1 Assessment of BGC descriptors

The comparative analysis of descriptors and classifiers on NPF-filtered and NPF-latest datasets is presented to measure the relevance of descriptors. Figure S1 presents the outcomes on the NPF filtered dataset that has 1003 instances and extracted features are 1191. Likewise, Figure S2 presents the outcomes on the NPF latest dataset containing 947 instances and 1157 extracted features.

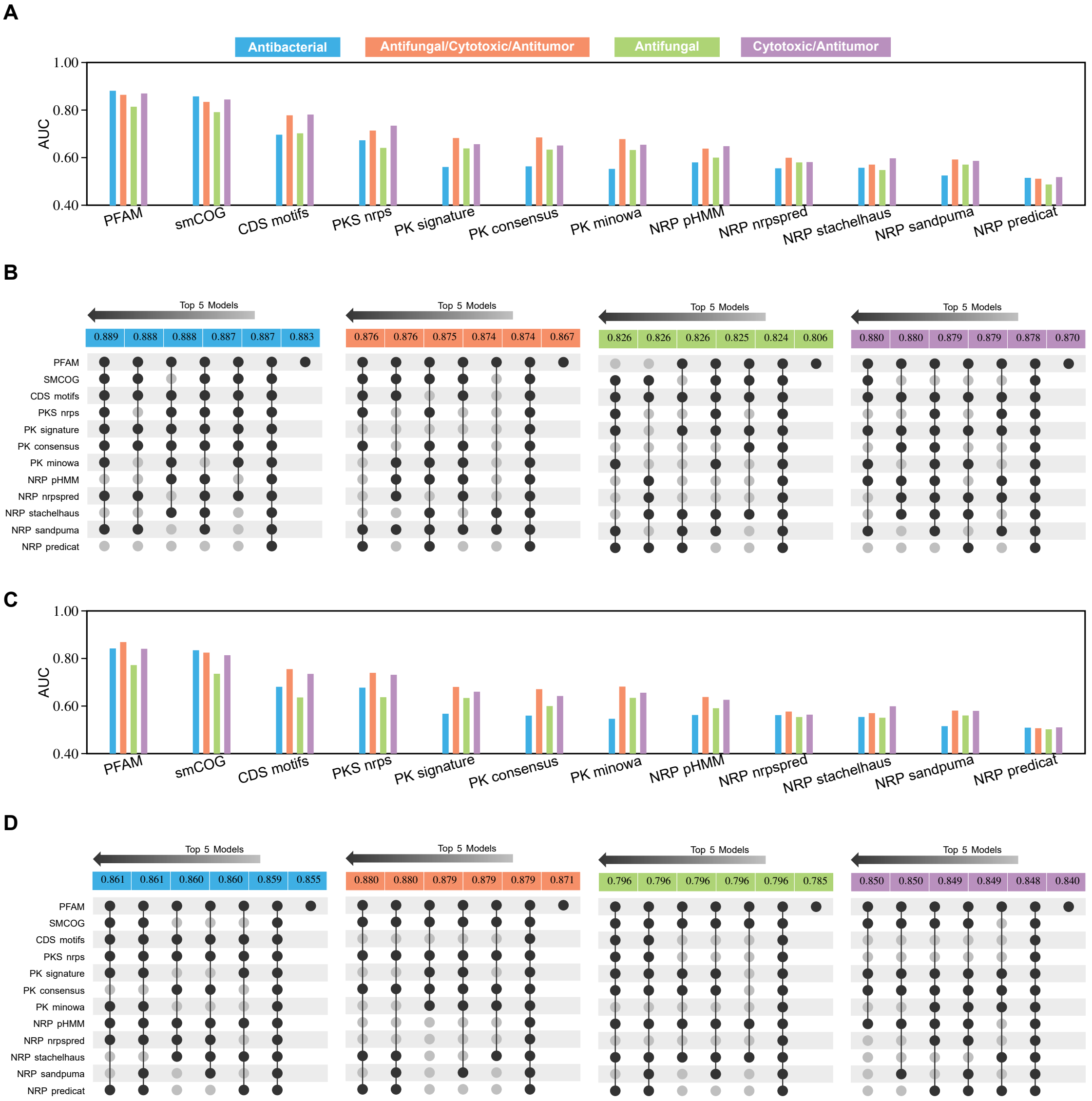


Figure S1: Performance analysis on NPP-filtered instances using ERTree and XGBoost classifiers. (A) Descriptor-wise performance and (B) performance of top 5 subsets of descriptors, in terms of AUC and their comparison with all descriptors as well as best descriptor (i.e., PFAM) for ERTree classifier. (C) Descriptor-wise performance and (D) performance of top 5 subsets of descriptors and their comparison with all descriptors as well as best descriptor for XGBoost classifier

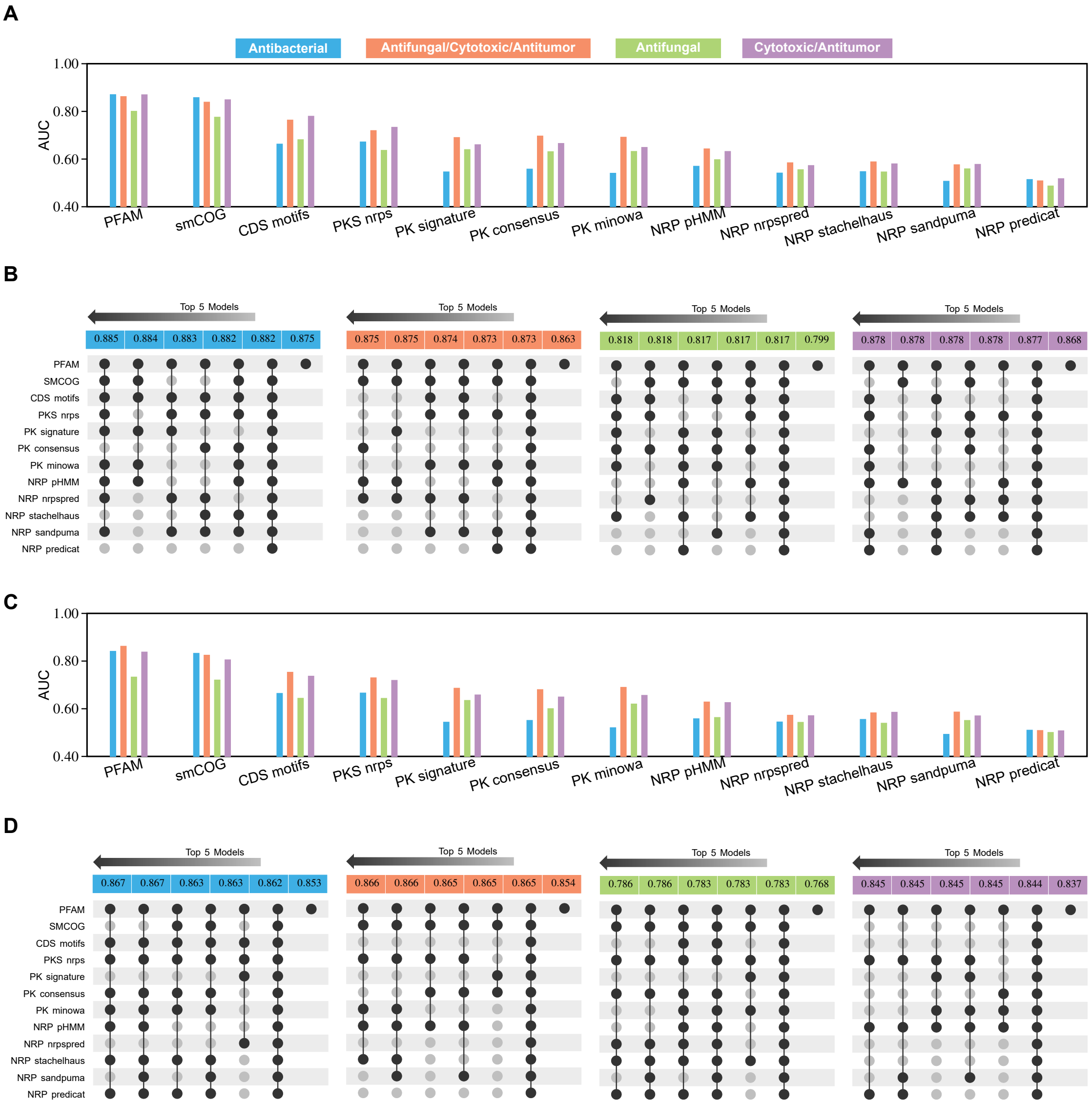


Figure S2: Performance analysis on NPP-latest instances using ExtraTree and XGBoost classifiers. (A) Descriptor-wise performance and (B) Performance of top 5 subsets of descriptors, in terms of AUC and their comparison with all descriptors as well as best descriptor (i.e., PFAM) for ERTree classifier. (C) Descriptor-wise performance and (D) performance of top 5 subsets of descriptors and their comparison with all descriptors as well as best descriptor for XGBoost classifier

#### 2 Problem of overfitting

Problem of under and over-fitting is common for models when underlying training sets are small. The models do not have enough information to capture the underlying patterns. Here, we measured the overfitting by assessing the NPF model on the 348 MIBiG instances which are not part of NPF-latest training set. The outcomes on these gold-standard instances are shown in Figure S3 which indicate high over-fitting for ERTree and XGBoost classifiers. For the antibacterial, antifungal, and cytotoxic/antitumor bioactivities, the difference is approx. 20% on ERTree classifier. Similarlity, the differences are approx 20% for the antibacterial, and cytotoxic/antitumor bioactivities and 10% for antifungal bioactivity on XGBoost classifier.

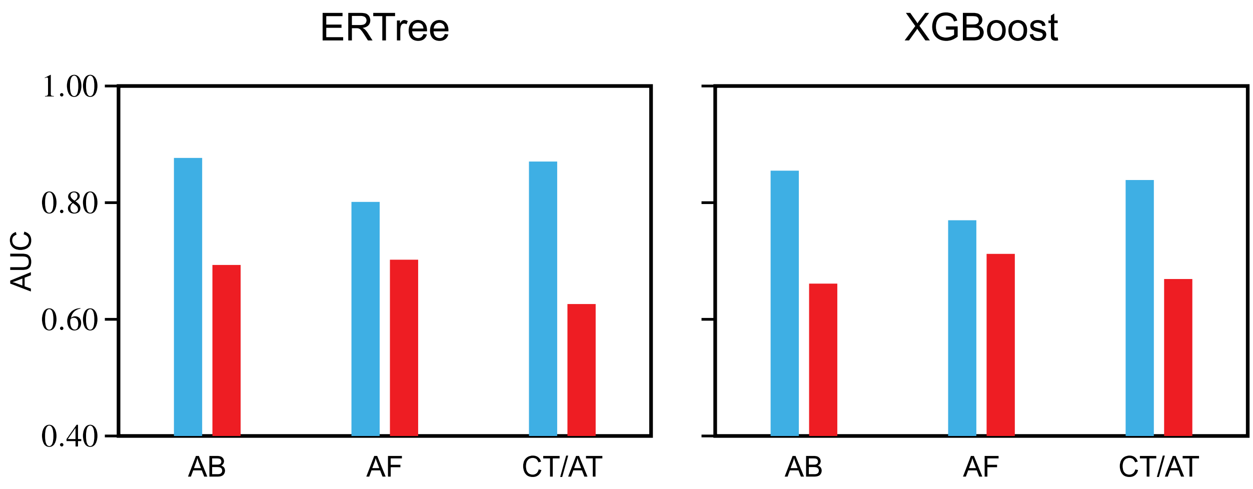


Figure S3: AUC outcomes for different bioactivity classes on NPF training (blue) and MIBiG test (red) sets. AB denotes antibacterial, AF denotes antifungal, and CT/AT denotes antitumor or cytotoxic

### 3 Assessment of PFAM and sequence descriptors for neural network model building

Figures S4-S8 present the outcomes of combinations of PFAM and sequence descriptors in terms of BCE training and validation loss. Several combinations of sequence descriptors, nucledotide profiles (NP) and protien profiles (PP), with PFAM are tried which include

- **NP**: 4, 6, & 8
- **PP**: 1 & 2
- **NP**: 4, 6, & 8 + **PP**: 1 & 2
- **NP**: 6, & 8 + **PP**: 1 & 2
- **NP**: 8 + **PP**: 1 & 2
- **NP**: 4, 6, & 8 + **PP**: 2
- **NP**: 6, & 8 + **PP**: 2

Figure S4 presents the outcomes of combination of PFAM and all nucleotide profiles only whereas Figure S5 presents the outcomes of combination of PFAM and all peptide profiles only. Figure S6 presents the outcomes of combination of PFAM and all nucleotide as well as peptide profiles. Figure S7 presents the outcomes of combination of PFAM and 6 & 8-mer nucleotide as well as all peptide profiles. Similarlily, Figure S8 presents the outcomes of combination of PFAM and 8-mer nucleotide as well as all peptide profiles. Figure S9 presents the outcomes of combination of PFAM and all nucleotide profiles as well as 2-mer peptide profiles.

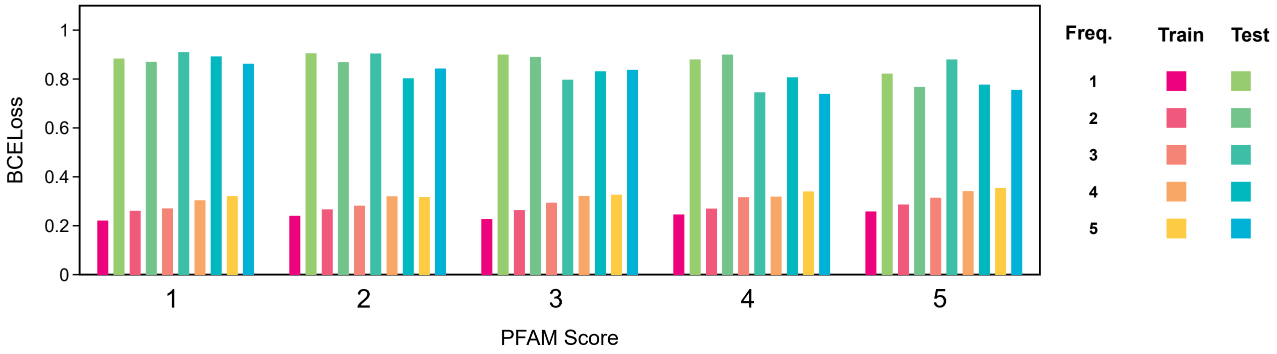


Figure S4: Performance with combination of PFAM and sequence descriptors (nucleotide profiles with 4, 6 and 8-mers)

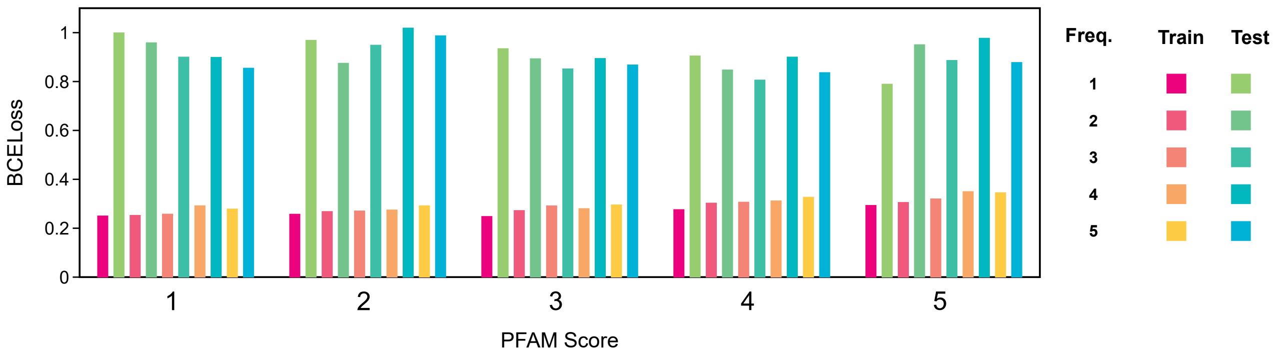


Figure S5: Performance with combination of PFAM and sequence descriptors (peptide profiles with 1 and 2-mers)

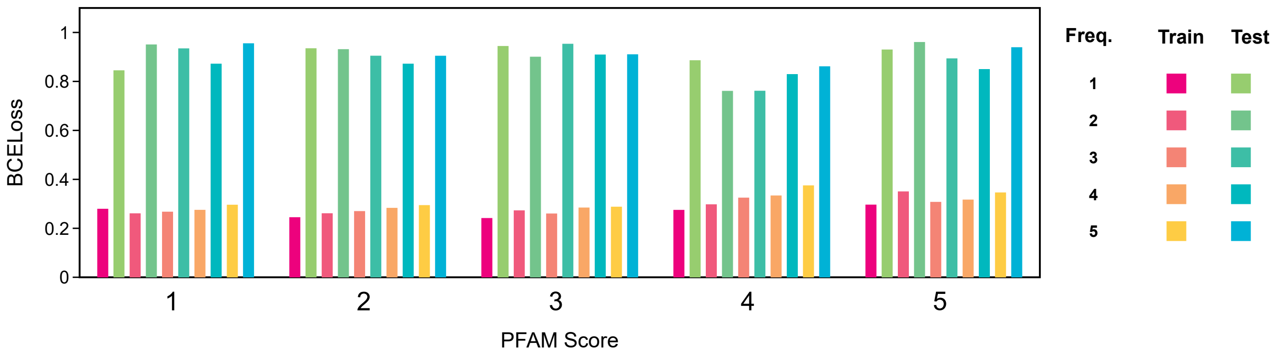


Figure S6: Performance with combination of PFAM and sequence descriptors (nucleotide profiles with 4, 6 and 8-mers and peptide profiles with 1 and 2-mers)

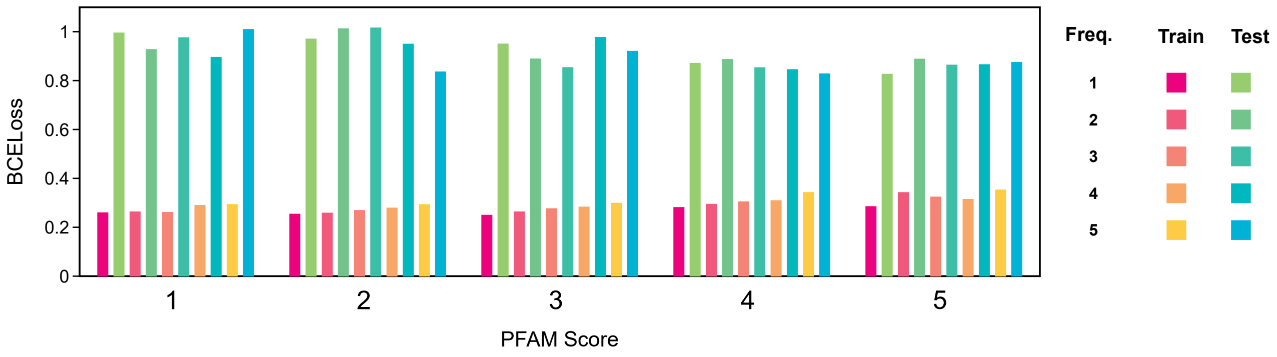


Figure S7: Performance with combination of PFAM and sequence descriptors (nucleotide profiles with 6 and 8-mers and peptide profiles with 1 and 2-mers)

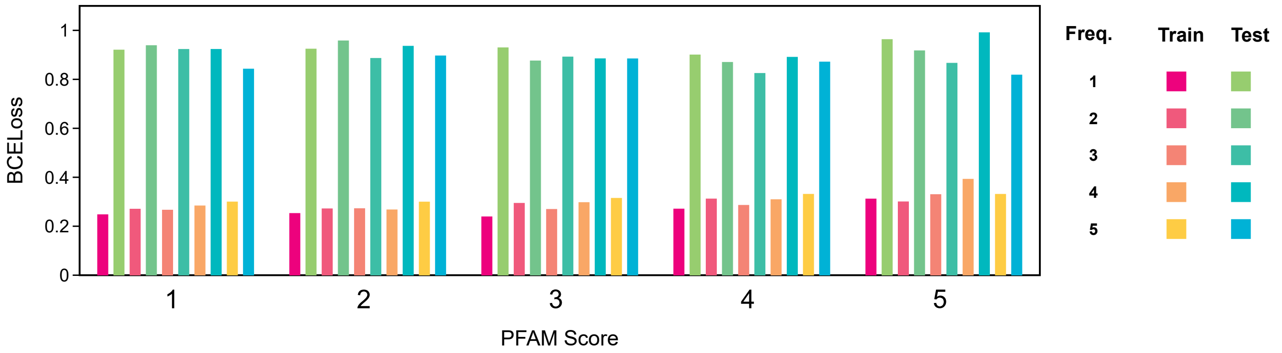


Figure S8: Performance with combination of PFAM and sequence descriptors (nucleotide profile with 8 mer and peptide profiles with 1 and 2-mers)

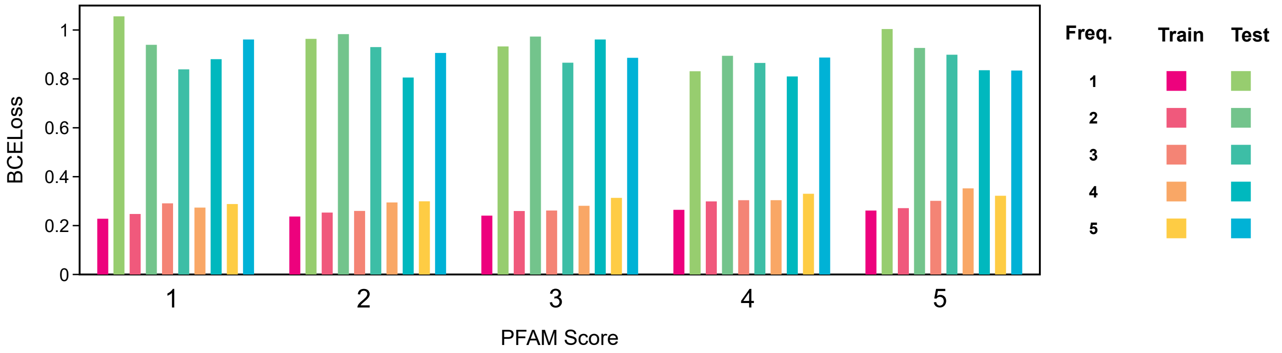


Figure S9: Performance with combination of PFAM and sequence descriptors (nucleotide profile with 4, 6 and 8-mers and peptide profiles with 2-mers)

### 4 Predictions on 635 unannoatated MIBiG BGCs

Figure S10 represent the the prediction outcome of NPBdetect on 635 unannotated MIBiG BGCs. Figure shows the highest prediction annotation count for antibacterial bioactivity with significant numbers of BGCs displaying multiple bioactivities too.

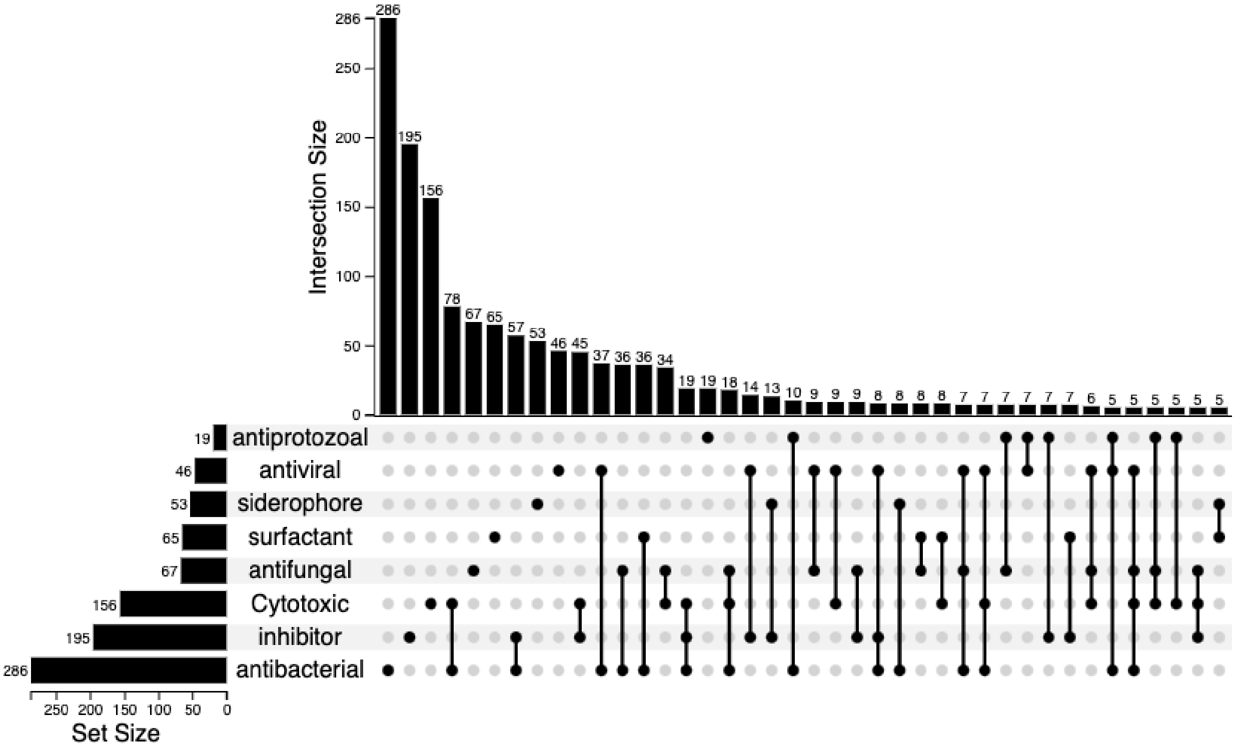


Figure S10: Distribution of the predicted Bioactivities for 635 unannotated MIBiG BGCs
